# Ultra-efficient, unified discovery from microbial sequencing with SPLASH and precise statistical assembly

**DOI:** 10.1101/2024.01.18.576133

**Authors:** George Henderson, Adam Gudys, Tavor Baharav, Punit Sundaramurthy, Marek Kokot, Peter L. Wang, Sebastian Deorowicz, Allison F. Carey, Julia Salzman

## Abstract

Bacteria comprise > 12% of Earth’s biomass and profoundly impact human and planetary health.^1^ Many key biological functions of microbes, and functions differentiating strains, are conferred or modified by genome plasticity including mobilization of genetic elements, phage integration, and CRISPR arrays. Characterizing each of these processes is time-consuming and requires custom bioinformatic workflows ill-suited to enable discovery of new sources of genetic diversity or to uncover which elements are active. Further, strain typing of bacterial species and approaches to discriminate sub-populations remain time-consuming and resource intensive. Here, we show that SPLASH, our published approach for reference-free discovery and analysis directly from raw reads, and an improved statistical assembly algorithm, compactors, unify diverse tasks in microbial sequence analysis: discovering new mobile elements and CRISPR arrays missing from any reference, and generating rapid, metadata-free strain typing of diverse bacteria. SPLASH and compactors together constitute a new general discovery tool for biological discovery in the microbial world.

## Introduction

Microbes possess an exceptional degree of genetic diversity within species and strains.^2^ This genomic plasticity can drive differences in phenotypes, such as virulence and antibiotic resistance. Yet, only a fraction of the sequence diversity these processes generate can be captured by reference genomes. Even mapping to a reference with a single allele difference can be challenging and confound inference.^3^ Beyond limiting biological discovery, this approach is resource intensive, and this problem is exacerbated in highly polymorphic populations of microbes. The limitations of assembly and alignment hamper discovery of novel mechanisms that diversify genomes, as well as efforts to characterize strains and their phylogenetic relationships.

A new approach is needed, especially as the scale of microbial data collection grows.^4^ We recently introduced SPLASH, an approach that directly analyzes raw sequencing data to detect a signature of sample-specific sequence variation.^5^ This variation is detected in variable sequences (targets) proximate to a short constant sequence (anchor), where every *k*-mer in a sequencing dataset is considered as an anchor (Figure 1A). We have previously shown SPLASH to unify detection of myriad forms of sequence variation and perform inference without any reference genome or metadata, such as knowledge of sample origin.^5^ By theoretical design, SPLASH should unify detection of structural variants, mobile elements and diversifying sequences in complex microbial populations. SPLASH is also ultra-efficient: it is many times faster than the alignment step in standard workflows; its speed is independent of genome size.^6^

**Figure 1.**
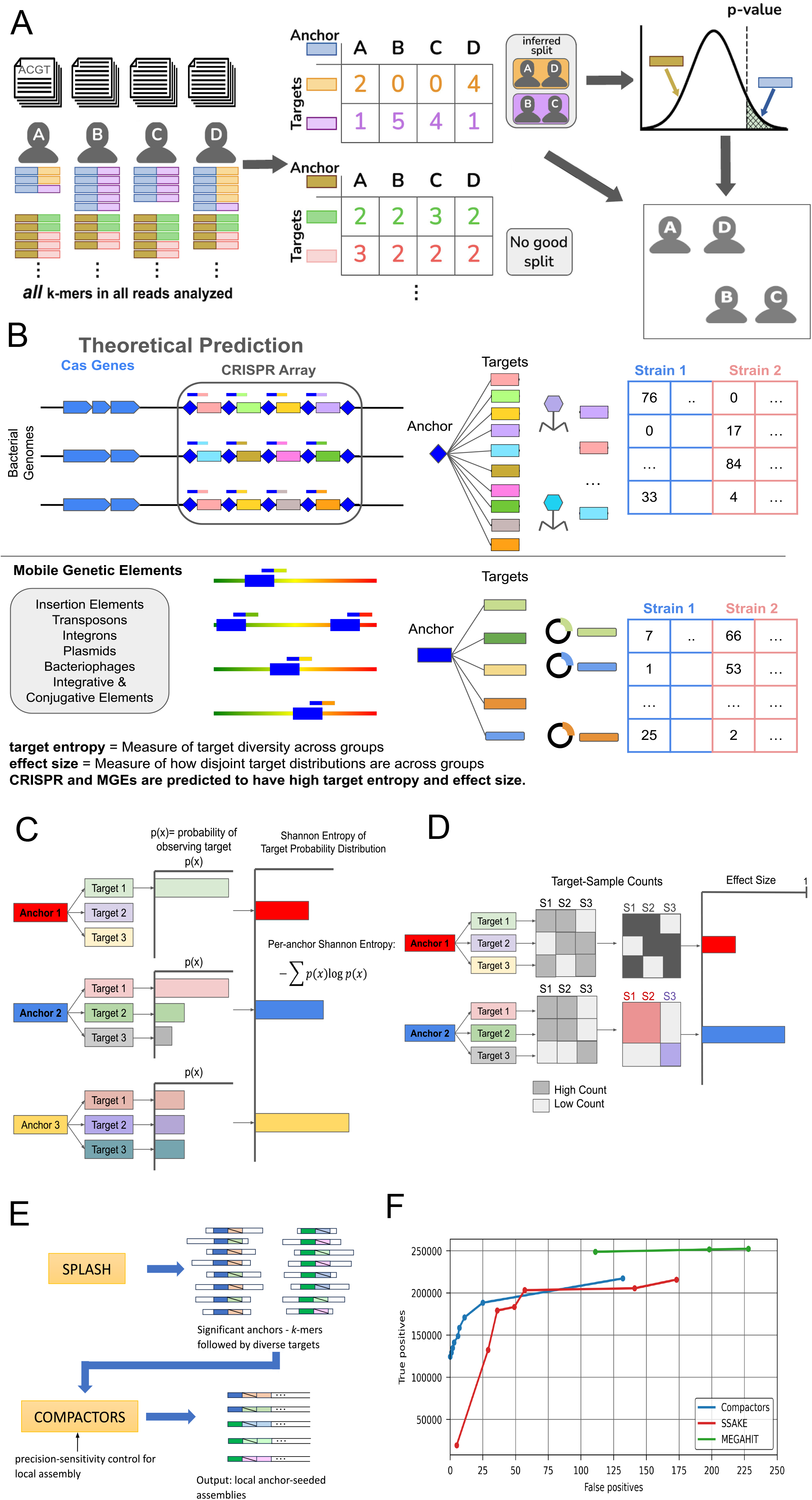
Compactors enable efficient de novo assembly; statistical metrics define CRISPR and mobile genetic elements. (A) SPLASH parses all *k*-mers in all sequencing reads. Each *k*-mer that exists in the reads is called an anchor, and the *k*-mer directly following the anchor is defined as the anchor’s target. Each anchor’s target counts across samples are collected in tables. On these tables, SPLASH performs a statistical test of whether an anchor’s target distribution signals sample-specific sequence variation. Anchors with low SPLASH p-values and high effect sizes capture biological sequence variation in their targets, including single-nucleotide polymorphisms, insertions and deletions. Aggregating this information across many anchors, SPLASH clustering groups samples by their distributional concordance across all such detected variation. (B) SPLASH anchors found in CRISPR repeat sequences or mobile genetic element (MGE) termini have diverse downstream targets corresponding to spacer sequences or integration loci. SPLASH statistically detects CRISPR-Cas systems and mobile genetic elements, including through target count variation across samples. (C) The distribution of target counts across all samples is used to measure the anchor’s Shannon entropy–the more uniform and diverse the targets, the higher the Shannon entropy (illustrated for a toy example with 3 targets). (D) Effect size measures and increases with the capacity of samples to be separated into two groups by their target distributions, which is possible when targets are very diverse. (E) Compactors improve local assembly of regions that are diverse across samples. Multiple algorithm parameters allow the user to control the tradeoff between the method’s precision and sensitivity. (F) Compactors improve control of true and false positives compared to SSAKE and MEGAHIT using the same protocol as in Suppl. Figure 2B.

We briefly outline why SPLASH is theoretically predicted to unify discovery in microbial sequencing data. SPLASH characterizes variation using *k*-mer pairs called anchors and targets (Figure 1A) (*k* = 27 by default but is adjustable). Every *k*-mer in the data is an anchor; each *k*-mer a fixed offset downstream (*R*, which may be zero) from a given anchor is one of its targets, always defined relative to an anchor. Anchors with more than one target can report on most sequence variations of interest: from changes at a single position to insertions, deletions and more. For this reason, SPLASH should discover active mobile genetic elements through conserved anchor sequences with highly diverse downstream targets varying by sample. The terminal ends of active MGEs should have high sequence diversity outside of the MGE; SPLASH will in principle call an MGE’s left and right termini as anchors, each having targets corresponding to loci where the MGE has integrated (Figure 1B). Conversely, mobile elements with no recent activity should have similar positions across strains; while their targets will be diverse, the distribution of target counts will not vary identically across samples. Thus, SPLASH would not be expected to detect ancestrally active mobile elements. SPLASH should also discover a host of other bacterial genomic features, from SNPs and indels to more rapidly evolving systems such as CRISPR.

For biological interpretation, a large sequence context around anchors identified by SPLASH is needed. Assembly is a traditional approach to this problem. Like alignment, assembly algorithms are rooted in a computer scientific past and, with few notable exceptions, have no statistical guarantees.^7^ Assembly algorithms have been shown empirically to struggle when generating contigs from complex samples such as in metagenomics and even assembly of repetitive regions of bacterial genomes.^8^ Moreover, assemblers have unknown false positive and false negative rates. To enable SPLASH for discovery in microbial genomes, we introduce a new statistical approach for assembly, significantly extending past work on the first, limited version of compactors.^9^ This new development extends SPLASH and enables a unified, hyper-efficient, reference- and metadata-free algorithm for microbial genomics.

We show that SPLASH discovers known and novel mechanisms that diversify genomes in both microbial isolate and metagenomic samples. We analyzed data from bacterial isolates of *E. coli*, *V. cholerae*, *M. tuberculosis*, *L. monocytogenes*, Groups A and B *Streptococcus*, and *S. pneumoniae,* including clinical samples and two studies of the human gut microbiome, one using dissolvable devices to sample the intestinal tract, another, the Inflammatory Bowel Disease Multi’omics Database (IBDMDB).^10–19^ We chose these datasets because they comprise well-characterized human pathogens and therefore provide a framework to assess how SPLASH compares to classical approaches to identifying bacterial strains and genetic features. We provide detailed study of the data, showing clear examples of novel and automatic discovery provided by SPLASH; processed outputs of species not discussed in detail will serve as resources for the community to explore.

A glimpse of SPLASH’s findings includes detection of integron arrays, a new CRISPR repeat and Cas-homolog missing from databases, and previously undescribed predicted miniature inverted-repeat transposable elements (MITE), a class of non-autonomous elements with a proposed role in gene expression. Further, the SPLASH algorithm and workflow performs *ab initio* phylogenetic discrimination including in *Mycobacterium tuberculosis* (*M. tb*), and unbiased re-discovery of a temporal shift in ICE element composition in clinical *Vibrio cholerae* samples. These disparate biological findings represent a snapshot of the potential sequence variation missing from reference genomes that are difficult or impossible to achieve with today’s assembly, alignment and bioinformatics pipelines.

## SPLASH and statistical *de novo* assembly (compactors) provide rapid analysis of diverse microbial studies

All analyzed data was downloaded from the NCBI SRA. The intestinal sampling study, the IBDMDB inflammatory bowel study, and the GWAS of *S. pyogenes* had the highest sum read count (respectively 14,587,329,831, 4,798,719,487 and 2,616,113,821 reads) and compressed size (resp. 824.1, 363.8 and 223.0 Gb).^14,18,19^ SPLASH runtime on each of these datasets was only 781.3, 332.5 and 132.6 minutes, respectively, roughly linear to sum FASTQ compressed size (Suppl. File 1, Suppl. Figure 1B). To interpret anchors called by SPLASH, we considered using currently available seed-based assembly packages. However, current *de novo* assemblers from short read data do not offer closed form statistical analysis of their rate of generating false positives. Further, they are resource intensive and struggle with assembly when the target is a pool of highly diverse sequences (such as a CRISPR repeat array).^20^ These limitations in the state-of-the-art prompted us to develop an alternative method.

To offer an alternative, we engineered a new statistical approach to local seed based assembly significantly improving our previous approach, called compactors.^9^ At a high level, the novelty of our development is iterative extension on a statistical criterion: the assembly will proceed as long as the terminal and seeding *k*-mer can be statistically linked (Methods). The statistical procedure provides parameters to control sensitivity versus specificity as a function of sequencing error and the length of the assembly (Methods, Figure 1F, Suppl. Figure 2B). We evaluated compactors compared to state of the art tools using synthetic and real data.

First, we tested compactors’ ability to reconstruct long local assemblies with simulated reads from a random 25-kbp reference generated using ART with built-in error models for HiSeq (read length up to 150-bp) and MiSeq (read length greater than 150-bp) devices.^21^ We tested whether compactors could reconstruct the middle 5-kbp of the reference (10,000:15,000) by seeding the algorithm with a 27-base anchor at position 10,000 in the genome (changing the query fragment had no effect on the performance as long as it was sufficiently far from begin/end of the reference to obtain uniform read coverage). The length of the assembly reconstructed by compactors depended on the stringency of the probabilistic criterion for compactor extension (parameter *min_extender_specificity*) (Suppl. Figure 2A). For parameter values of .6 and lower, compactors reconstructed the entire 5000-bp fragment at 60x coverage independently of the read length. We compared performance of compactors to widely-used assembly algorithms: MEGAHIT and SSAKE.^22,23^ While SSAKE has an option to provide a set of seed sequences to the assembly process, it was not able to reconstruct the assembly when using the anchor as a seed. Therefore, a default mode which generates seeds from a read set was employed. MEGAHIT achieved similar effectiveness at three times lower coverage. SSAKE, however, was unable to reconstruct the entire assembly even at (∼100x) coverage.

We next tested compactors’ sensitivity and precision in real data using *Vibrio cholerae* isolates (SRR14297458-67). Every read from the isolate data was split in half, resulting in a twice-as-large set of synthetic reads, each read half the length of the originals. These synthetic reads were input to compactors and competing assemblers to assess their ability to reconstruct sequences present in the original reads. A resulting compactor (or an assembly fragment starting from an anchor) was counted as a true positive if it appeared in the original reads more than once (a single occurrence likely corresponds to a sequencing artifact). We used the 500,000 highest effect size anchors called by SPLASH as seeds to compactors and SSAKE (MEGAHIT does not use seeds).

A desired property of any algorithm (or statistical test) is the existence of a parameter regime where only true positives are called; as the parameter is relaxed, the ratio of true positives to all calls decreases. Compactors has several parameters that control TP-FP trade off, producing the strictly monotonic trajectories we expect (Suppl. Figure 2B) with *lower_bound* (the minimum number of reads supporting a compactor) and previously described *min_extender_specificity* having the largest effect on precision/sensitivity balance. However, we could not find such regimes for SSAKE or MEGAHIT. In SSAKE, the minimum base ratio (-r) gave the most stable control of the true/false positives. SSAKE had markedly lower sensitivity than compactors in regimes with very low false positive rates (*lower-bound* parameter was used for controlling the latter algorithm) (Figure 1F).

For instance, compactors at 0 FP rate identified more than 140,000 true positives while SSAKE at the most precise configuration with 5 false positives had less than 20,000 true positive calls. Controlling the TP-FP trade off in MEGAHIT was harder to predict. By a grid search in a parameter space, we found regimes characterized by the largest sensitivity and largest precision. The sensitivity of MEGAHIT was mildly affected by the parameters and larger than that of competitors in their most sensitive variants (∼250,000 vs ∼220,000 calls). On the other hand, the FP rate exceeded both compactors and SSAKE, with more than 100 false positives in this regime (Figure 1F).

Beyond precision, compactors is significantly faster than MEGAHIT and SSAKE, enabling rapid and high-throughput data processing. Compactors, MEGAHIT and SSAKE in their most sensitive configurations (rightmost points at Figure 1F) completed analyzes in, respectively, 0m28s, 4m14s and 25m11s requiring 1.9 Gb, 1.9 Gb and 21.3 Gb of RAM (compactors and MEGAHIT, as multithreaded tools, were run with 16 threads). Compactors was approximately 8 times faster than MEGAHIT and almost 50 times faster than SSAKE; compactors’ memory-efficiency and statistical properties, including sensitivity and specificity, provide clear advantages for large scale analysis of complex samples.

## SPLASH rediscovers known CRISPR loci are predicts new ones

We next tested if compactors could predict new diversifying mechanisms in bacteria. To this end, we tested if SPLASH can rediscover CRISPR, one of the best studied immune systems in prokaryotes and well-known to generate high diversity in the spacer sequences–flanked by CRISPR repeats–in genomes harboring them.^24^ First, we assessed whether SPLASH identifies anchors at the “edge” of a larger sequence identifying a genome-diversifying mechanism. If so, compactors could illuminate the mechanism by providing sequence beyond the anchor and its targets. Under mild conditions, SPLASH should theoretically detect anchors in CRISPR repeats on purely statistical criteria: the fixed repeat sequence should be a SPLASH anchor, and the following set of variable spacers should be targets (Figure 1C). Active arrays should have great spacer diversity varying by sample, leading to high effect sizes (Figure 1D). As CRISPR arrays contain multitudes of self-dissimilar spacers, the distribution of their counts, if recorded in targets, will generate high Shannon entropy (Figure 1C). Further, because they are self-dissimilar, spacers’ joint Hamming distances should be high.

We tested if SPLASH had rediscovered CRISPR repeats among its targets as follows: For each of the analyzed datasets, the most entropic anchors having effect sizes of at least 0.9 were selected and mapped to CRISPRCasdb spacers and direct repeats as well as Rfam records reporting CRISPR elements (Table 1, Methods, Supplemental Methods, Figure 2B). *E. coli*, *L. monocytogenes*, *V. cholerae*, and Group A *Streptococcus* isolate sequencing, and IBDMDB, had more than 10 such anchors aligning to curated CRISPR repeat databases (Tables 1a-e). We used the Kolmogorov-Smirnov (KS) test to test if the Shannon entropy of anchors’ target distributions differed between anchors aligning to known CRISPR repeats and those aligning to other mobile elements (ICE) and all other sequences (Methods, Figure 2A). All p-values were significant (maximum and minimum p-values of 3.4e-07 in *Listeria* and 3.8e-24 in *E. coli*), establishing reproducible, statistically significant distributional differences between anchors aligning to CRISPR repeats and those that did not (Figure 2A). Anchors aligning to categories with high flanking diversity, such as iterative and conjugative elements (ICE) had less or no enrichment (File 2, Figure 2A, Supplemental Methods). Further, the majority of CRISPR-aligning anchors were in the 90th percentile of target entropy even within restrictive sets of anchors with the highest effect sizes (Methods): in IBDMDB, 31/58 CRISPR-aligned anchors were in the 90th percentile of entropy; 26/68 in *Listeria*; 37/67 in *V. cholerae*; 54/100 in *E. coli*, and 19/37 in Group A *Streptococcus* (Southon et al. 2020) (Table 2). The final measure distinguishing anchors aligning to CRISPR repeats was joint Hamming distance: the count-weighted average target Hamming distances were consistently higher in CRISPR-aligned anchors than others with differences between sets ranging from 3.2 to 5.7 (corresponding to ∼ 4.3 to 7.5 SNVs) in the datasets investigated (Table 2, Figure 2B). The statistical discrimination of CRISPR-aligning anchors raises the possibility that anchors having great diversity in target sequence composition and number may constitute discoveries of sequence-diversifying mechanisms similar to CRISPR.

**Figure 2.**
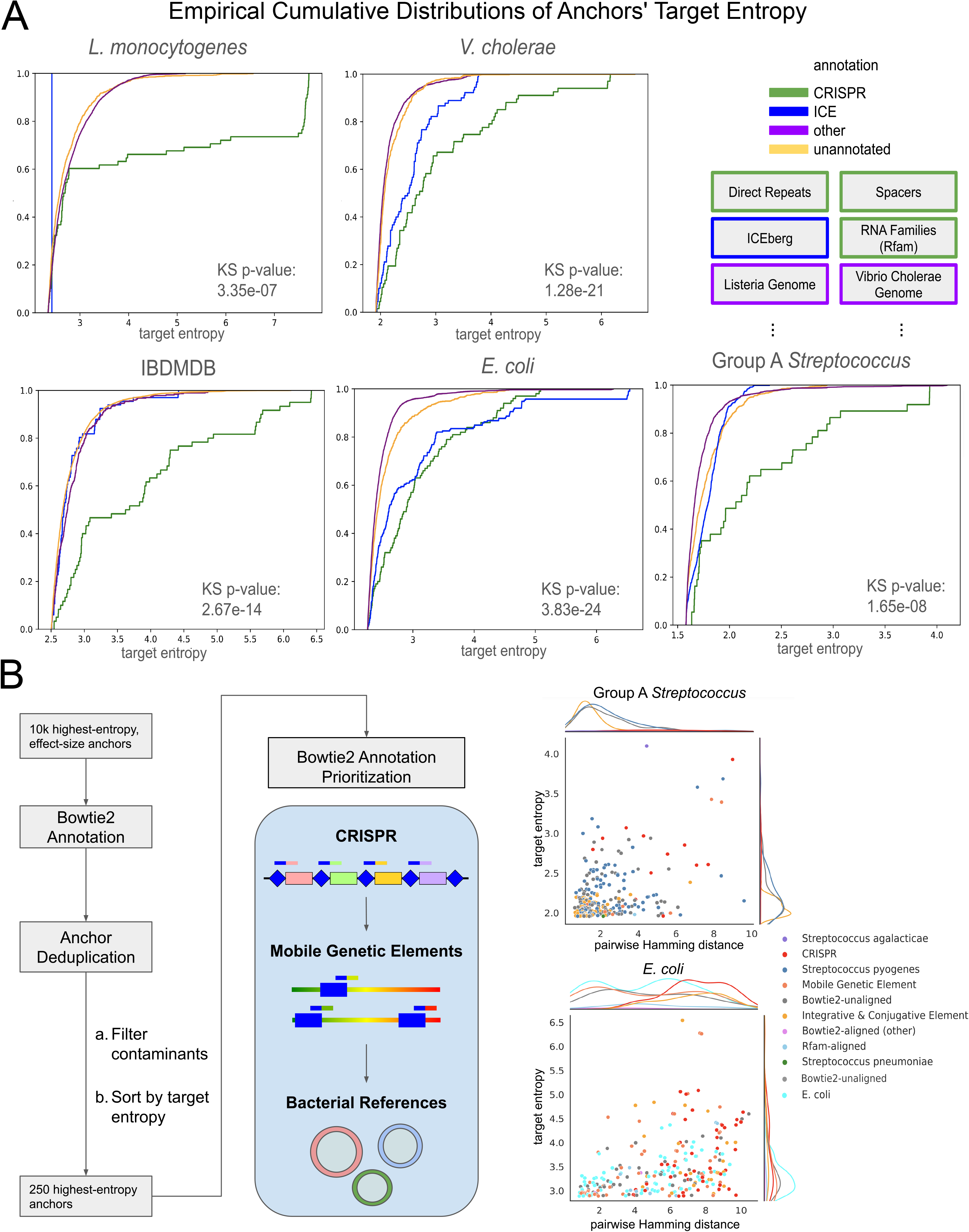
SPLASH rediscovers CRISPR and illuminates a novel CRISPR homolog. (A) SPLASH rediscovers active CRISPR systems in anchors having high target entropy. ECDFs are shown of anchors’ target entropy, where anchors are classified through Bowtie2 alignment to Rfam, ICE elements, CRISPR sequences, and bacterial genome assemblies (Supplemental Methods). The Kolmogorov-Smirnov (KS) test compares the distribution of CRISPR-annotated anchors to anchors not annotated as CRISPR; all KS p-values are highly significant (Methods). (B) A flowchart depicts Bowtie2 annotation and selection of the 250 highest-entropy and high effect size anchors. The hierarchy of annotation categories for anchors with multiple alignments is shown: CRISPR, for example, overrides anchors’ categorization as MGE or bacterial genomic sequence. In Group A *Streptococcus* (Southon et al. 2020) and *E. coli*, anchors categorized as CRISPR via Bowtie2 are shown to have the highest entropy and pairwise Hamming distance among targets. Anchors are processed to remove offsets (Methods) and those Bowtie2-aligning to contaminant databases (Supplemental Methods).

**Tables 1a-k:** Highest-entropy anchors subject to entropy and effect size filters. Anchors having effect size above 0.9, in the top 10,000 by entropy, and Bowtie2-unaligned to UniVec and Illumina adapters are reported with their Bowtie2 alignments to bacterial species and mobile and repetitive elements (Supplemental Methods). Tables sublineated as follows: (a) *Listeria* (Maury et al. 2016); (b) *V. cholerae* (LeGault et al. 2021); (c) IBDMDB; (d) *E. coli* (GenomeTrakr); (e) Group A *Streptococcus* (Southon et al. 2020); (f) Capsule Intestinal Profiling (Shalon et al. 2023); (g) Group B *Streptococcus* (Chaguza et al. 2022); (h) Group A *Streptococcus* (Kachroo et al. 2019); (i) *Streptococcus pneumoniae* (Chaguze et al. 2020); (j) *M. tuberculosis*, lineage 2 (CRyPTIC); (k) *M. Tuberculosis*, lineage 3 (CRyPTIC).

**Table 2:** Distributional differences in effect size, entropy, and pairwise Hamming distance. For each of Tables 1a-e, anchors are defined as CRISPR-aligned if Bowtie2-aligned to direct repeats and spacers, or to an Rfam taxonomy reporting CRISPR (Supplemental Methods). Average pairwise Hamming distance, target entropy, and effect size are reported separately for CRISPR-aligned anchors and all others.

## Discovery of a CRISPR repeat array absent from reference databases

Rediscovery of CRISPR on purely statistical metrics suggested that anchors with statistical scores similar to CRISPR repeats and missing from available references may represent new discoveries. We searched for anchors having high effect sizes and the highest entropy, further restricting to those absent from NCBI nucleotide databases (Methods, Table 3, Suppl. File 2A). The highest-entropy anchors had poor alignment rates, with 395 of the 4,261 anchors failing alignment, with no BLASTN hits against the nr database (Suppl. Figure 3A). For these 395 anchors, compactors were generated to investigate the sequence diversification driving the anchors’ high entropy (Suppl. File 2B).

**Table 3:** Highest-entropy anchors subject to further postprocessing. Anchors, subject to having entropy > 3, from select datasets in Table 1 chosen to support replicable analysis.

We tested if any of the highest entropy compactors could represent novel CRISPR arrays. In the top 10 anchors by entropy, we identified one anchor (entropy 5.1) whose compactors presented a repeat-spacer motif consistent with being a CRISPR array (Suppl. File 2C). The anchor was found in a 36-bp subsequence repeating at regular intervals in its compactors; distinct sequences of constant length interspace the repeats. Protein translations of the 23 compactors failed alignment to NCBI protein databases (Methods). Three compactors aligned to NCBI-reported assemblies annotated as phage, with e-values of 1e-04 (BK040081.1), 0.016 (OP031041.1), and 0.005 (BK018822.1) (Suppl. File 2D). Alignments covered compactor subsequences interspacing the repeated element we had identified, where they had 97-100% identity (Suppl. File 2D). CRISPR spacer sequences represent fragments of phage genomic sequence; thus, evidence of the spacer sequences in assemblies annotated as phage suggested we had captured an unannotated CRISPR array.

We hypothesized that the unannotated repeat array corresponded to a CRISPR-Cas system or to an undescribed system. As SPLASH anchor-targets and compactors were missing from NCBI databases, we queried Pebblescout–a tool to rapidly screen all NCBI-hosted metagenomics datasets–for studies reporting the repetitive 36mer in long-read sequencing (Supplemental Methods).^25^ Long reads enabled us to scan beyond the repeat array for evidence of a CRISPR-Cas system nearby. From 2 metagenomic studies, Pebblcout identified 4 FASTQs reporting PacBio and Nanopore long-read sequencing, containing 11,102,962 total reads (Supplemental Methods, Suppl. File 2F). Only 9 of these 11 million reads contained the 36mer, likely due to differential microbial abundance in metagenomic sampling. To evaluate homology to Cas and to profile the array’s genomic context, reads’ protein translations were aligned to NCBI databases (Methods, Suppl. File 2G). All reads aligned to one of Cas1, Cas2, or Cas9 with e-values ranging from 2e-125 to 1e-14 (Supp. File 2G). This result confirms homology to Cas; however, amino-acid identity between translated nucleotide sequences and the database proteins was as low as 412/1149 (35.8%) and 244/615 (39.7%) residues, indicating the proteins are substantially diverged (Suppl. Figure 3B, Suppl. File 2G). Across the 9 reads, only 1576 amino acids of the 3762 total covered by alignment had identity with the target protein sequences; individual reads had average percent identity of 49.8% to the aligned proteins.

Additional to alignment to Cas, A/G-specific adenine glycolase was hit, consistent with a requirement for DNA repair in regulating CRISPR-Cas mediated genome modification (Suppl. File 2G). *In silico* translation of long reads followed by Pfam alignment–a procedure we call “Pfam-analysis”–confirmed the presence of Cas proteins in 3 reads and provided further insights into other domains located close to the array: they include annotated as ATP-dependent helicase, RNA-binding and related to transposition (Methods, Suppl. File 2H).^26,27^ Together, this suggests that the new repeat array may house novel Cas-family proteins.

## *De novo* identification of integron arrays in clinical strains of *Vibrio cholerae*

CRISPR is only one of many known arrays in prokaryotes characterized by rapid sequence diversification. To investigate genomic sequence diversification beyond CRISPR, we focused on a study of hundreds of isolates of *Vibrio cholerae,* a water-borne pathogen endemic to southeast Asia that can cause severe diarrheal disease (PRJNA723557). We first identified anchors where target abundance predicted collection time and/or source (water versus stool) using general linear regression (Methods, Table 4).^28^ To guide interpretation, we analyzed anchors mapping to the *V. cholerae* genome (Supplemental Methods). Among these, the 5 highest-entropy anchors perfectly aligned near the 5’ and 3’ ends of a reported *V. cholerae* superintegron repeat: a 124-bp extragenic repeat: this array is thought to have dynamic length and to host genes in the sequences intervening repeats.^29^ Cassette genes are actively inserted and transcribed in the superintegron array, which has this name because, compared to other species’ integron arrays, in *V. cholerae* it can contain hundreds of cassette genes.^30,31^ Due to its repetitive nature, this superintegron has been difficult to characterize by traditional methods.

**Table 4:** Consolidated results of GLM run on anchor-target counts in samples. Result of glmnet multinomial regression for categories including collection year, country, and tissue, in Group A *Streptococcus* (Kachroo et al. 2019) and (Southon et al. 2020), Capsule Intestinal Profiling (Shalon et al. 2023), IBDMBD_PRJNA398089, *V. cholerae* (LeGault et al. 2021), *Streptococcus pneumoniae* (Chaguze et al. 2020), and *Listeria* (Maury et al. 2016) (Methods).

To illustrate SPLASH’s discovery power, we further investigated one anchor with 100% identity to within 1 base-pair of the 3’ end of the reported *V. cholerae* superintegron repeat, (entropy 6.75, SPLASH-called in 282/283 samples) (Suppl. File 3, Figure 3C). Targets of this anchor strongly predicted sample source: the 4th most-abundant target had the highest odds ratio (GLM coefficient 1.475) for stool versus water. The 1st and 4th most-observed anchor-targets had high counts in stool, appearing in all samples and being the most highly enriched targets in stool (10743/279143, 8156/279143 counts for anchor-target 1 and anchor-target 4, respectively; Figure 3B), indicating the cassettes associated with these targets are pervasive and enriched in *V. cholerae* found in stool. These results suggest association with adaptations of *V. cholerae* to human hosts.

**Figure 3.**
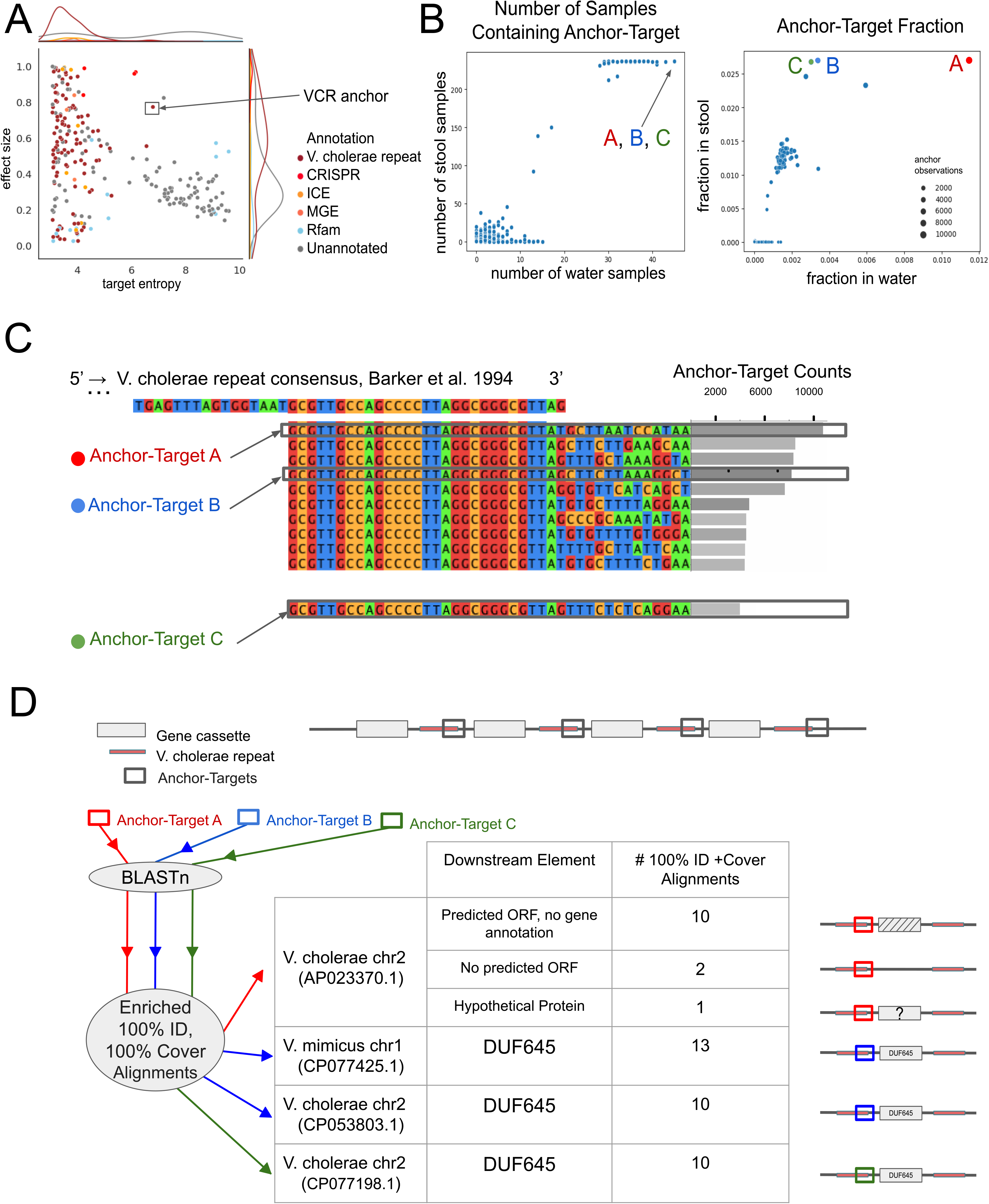
De novo detection and insight in the V. cholerae superintegron. (A) Bowtie2-unannotated and Rfam-aligned anchors have the highest entropy. Two anchors have high entropy and effect sizes: one fails Bowtie2 alignment and BLASTN-aligns 40 nucleotides upstream of hypothetical protein QCO92807.1 in uncultured bacterium clone contig MK411236.1; the other Bowtie2-aligns to *V. cholerae*, matching the intergenic repeat of the *V. cholerae* superintegron. (B) Anchor-targets A, B, and C were the only 3/6311 anchor-targets detected in all samples (A, B, C represent anchor-targets 1, 4, and 30, respectively). A, B, and C were the most commonly-observed in samples annotated as stool and among the most commonly-observed in samples annotated as water, of the 792 targets observed at least 5 times. (C) Anchor-targets A, B, and C and their counts, with multiple sequence alignment to the *V. cholerae* superintegron repeat consensus sequence. (D) Anchor-targets are expected to align between two gene cassettes, as the anchor corresponds to the intergenic repeat. BLASTN mappings are described in terms of the adjacent ORFs, communicating the lack of functional annotation for cassettes associated with anchor-targets A, B, and C.

We next used compactors to investigate cassette genes downstream of anchor-targets 1 and 4. We hypothesized that the enrichment in counts of anchor-targets 1 and 4 was potentially consistent with a regulatory mechanism inducing cassette gene amplification; we investigated this using Pfam analysis of compactors (Methods). 5 of 42 compactors that extended anchor-target 1 mapped to DUF3709, a domain reported to be proximal to reverse transcriptases, and 33 had no hits to Pfam.^32^ This finding–bypassing any reference alignment– is further supported by the anchor-target mapping to a *V. cholerae* assembly upstream of a gene including DUF3709 (Suppl. File 3I).

Eleven distinct compactors had Pfam domain hits to ParE_toxin, a positive control known to regulate chromosome stability in *Vibrio* (93 compactors, average e-value 1.7e-05) (Suppl. File 3I).^33^ Ten distinct compactors, including 6 compactors stemming from anchor-target 4, which had strong association with collection from stool, mapped to DUF645, a second domain reported to neighbor a reverse transcriptase (Suppl. File 3I).^34^ Further analysis of anchor-targets 4 in annotated *Vibrio* assemblies also suggests a diversity of genes with DUF645 domains at multiple copy number (Suppl. Methods). Other domains that appeared in five or more distinct compactors include several domains associated with (anti)toxin function including PhdYeFM_antitox (35 compactors, average e-value 2.7e-04) and YafQ_toxin (23 compactors, average e-value 9.5e-12), as well as domains NTP_transf_6, a nucleotidyltransferase superfamily member of unknown function, (45 compactors, average e-value 1.6e-09) and a set of domains annotated as acetyltransferases, including FR47 (Suppl. File 3I). The SPLASH anchor-targets associated with DUF645 guiding this analysis were present in all samples and had the highest counts in stool; with the absence of an annotated function, this domain is a compelling object of future study. These results demonstrate how SPLASH and compactors can provide a path forward for prioritizing domains–with or without Pfam annotations–for further study.

## Protein domain analysis of compactors identifies new mobile elements

We sought to test if SPLASH and compactors could identify new mobile elements or domains involved in mobilization, reasoning that a mobile element should have a highly diverse downstream sequence context, eg: be identified by SPLASH, but have a stereotyped upstream sequence (Figure 4A). To identify sequences with the latter quality, compactors were seeded with anchors’ reverse-complements. If a single compactor was generated, Pfam-analysis of that compactor was performed (Methods, Figure 4A, Suppl. File 4A, 4B). We then computed the 80th percentile entropy and effect size of anchors with compactors assigned to each domain that had at least 10 contributing anchors (Suppl. File 4D). Domains with the highest entropy and effect size were enriched for Cas proteins, transposases, and other domains known to be involved in mobilization of DNA (Figure 4B). Many other domains were also enriched, including several “Domains of Unknown Function” (DUFs) including DUF4368 and DUF6783, NDUFV3 and LRR_5, each surprising and suggesting a potential mismatch between the annotated domain and mechanism generating diversity. To test this, we computed the average query cover and percent identity when compactors were BLASTN-aligned against NCBI nucleotide databases (Methods, Figure 4C).

**Figure 4.**
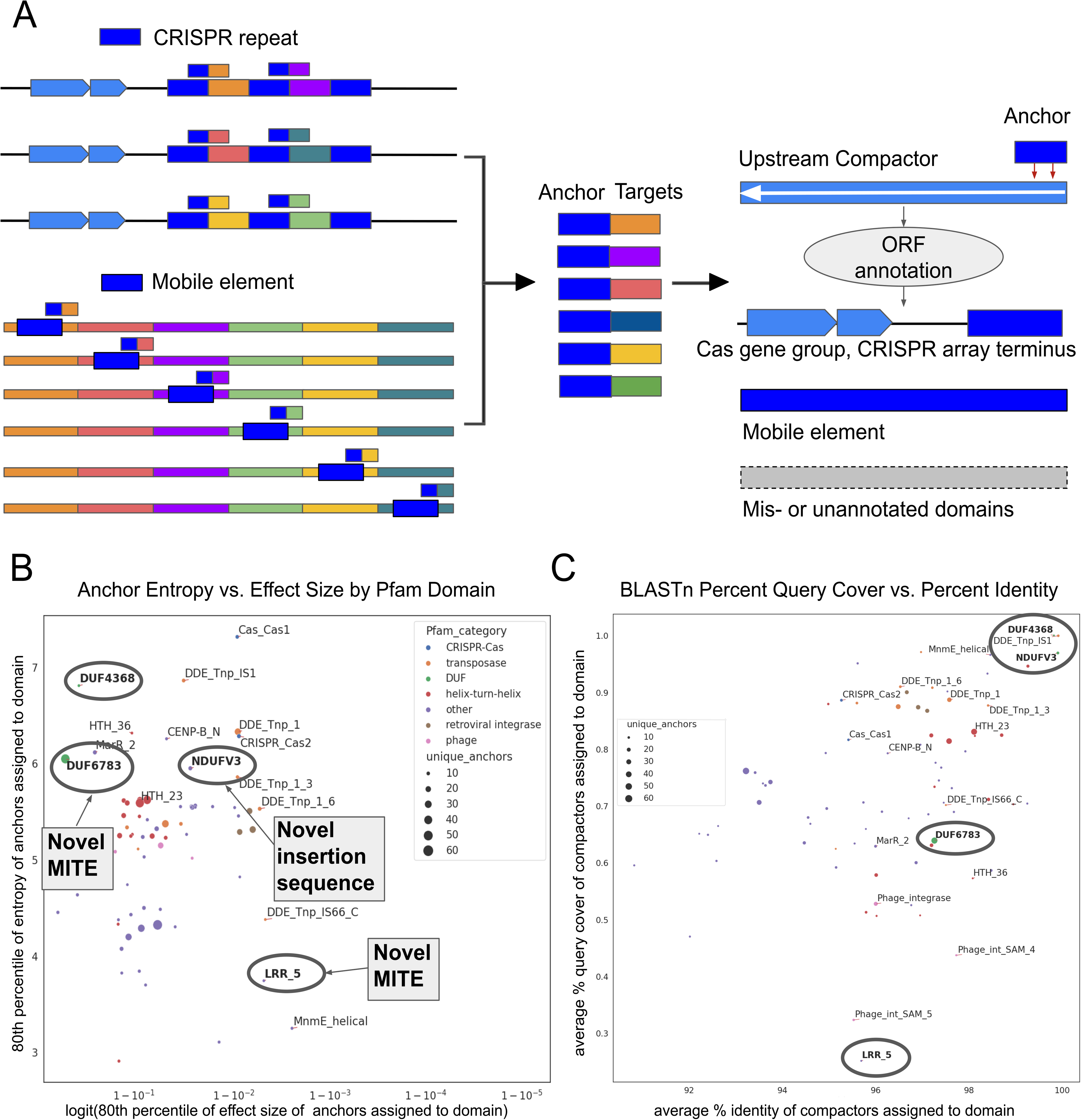
Compactors and SPLASH statistics prioritize novel transposable elements. (A) SPLASH captures CRISPR repeats and mobile elements through anchors having highly diverse targets. While MGEs are characterized by high target diversity downstream of the MGE end, the upstream sequence should be stable. Thus, local assembly opposite the direction of diverse targets is predicted to capture the diversifying mechanism. (B) Compactors were generated upstream of SPLASH-called anchors and selected if only one underlying upstream sequence was generated. Transposases, CRISPR-Cas, and retroviral integrase domains were highly enriched in compactors whose anchors had high effect sizes and entropy; other domains, such as DUF4368 and DUF6783, had entropy matching transposase domains. (C) After BLASTN search result of compactors in (B), the average BLASTN percent query cover and identity of compactors Pfam-aligned to each domain. Low query cover in compactors aligning to Phage integrase was hypothesized to be due to the high rate of viral evolution.

We first investigated DUF4368; Pfam annotation states that it is associated with resolvase/recombinase domains. Recent work has shown it to be associated with multi-targeting large serine recombinases (ones with relatively loose target site specificity).^35^ BLASTN of DUF4368 compactors shows they are associated with integrative mobilizable elements (IMEs), which are related to ICEs; IMEs have a recombinase gene for integration/excision and a mobilization relaxase gene for conjugative transfer, but they lack most genes required for conjugation.^36^ Compactors’ BLAST reveals three IME groups with different associated genes.

The first group has a virulence-associated protein E gene; the second group has a tetracycline resistance gene; and the third group has multidrug transporter genes.

We next investigated DUF6783; BLAST showed that compactors Pfam-aligned to the domain were embedded in the coding region of a variety of genes, typically in *Eubacteriales* (e.g. *Blautia* and *Ruminococcus*) with a 162-nt core sequence with one open reading frame on each strand and inverted repeat symmetry. We found examples of the same host gene with and without insertion of the 162-nt sequence, indicating it is mobile. While the sequence is found within protein-coding genes, BLASTN shows the majority of occurrences are intergenic. These qualities match with those of miniature inverted repeat transposable elements (MITEs), which are thought to be non-autonomous elements reliant on a co-resident transposase, and can carry RNA or DNA motifs. We know of only one other example of a MITE that can insert in-frame into coding sequences, the NEMIS element in *Neisseria*; the NEMIS element has not been assigned a Pfam domain, unlike DUF6783.^37^ It should be noted that for DUF6783 we do not find target-site duplications as would be expected for insertion by a transposase. The consensus insertion site is TT|AG, and it often inserts precisely at a TAG stop codon.

BLASTN of compactors associated with high entropy and effect size anchors reported LRR_5, a leucine-rich-repeat protein, and Phage integrase domains as having the lowest summary BLASTN alignment quality. We investigated compactors Pfam-aligning to LRR_5 as those had the lowest average BLASTN percent query cover and e-value (25.11%, 1.29e-03) (Figure 4C, Suppl. File 4D). We investigated one anchor further: it appeared to have highly sample- and donor-specific compactors (Methods, Suppl. Figure 4, Suppl. File 5). The majority of anchors associated to this domain are intergenic, suggesting it may be a novel mobile element, and are not mapped near LRR_5, meaning that there may be an enrichment of the insertion near an LRR_5 domain or the association is incidental. The anchor is often found via BLASTN as part of a nearly palindromic ∼159-bp sequence present multiple times in different *Segatella copri* genomes, and thus has features of a MITE, though the sequence varies between examples and target-site duplications are not evident (target site often has additional palindromic nature). Our final investigation was of NDUFV3, a domain annotated as related to mitochondrial NADH-ubiquinone oxidoreductase flavoprotein 3. However, similar to the LRR_5 this Pfam domain assignment is likely incidental– a plurality but not majority of compactors contain the domain which is responsible for its call, but in fact, the compactors represent a novel insertion sequence of 1,419 nt, containing a transposase in the ISL3 family (compactors BLAST to the noncoding inverted repeat).

## SPLASH provides reproducible de novo and discovery of strain variation without phylogenetic modeling

In addition to discovery, SPLASH anchors together provide information on whether sample-level sequence variation has consistent patterns between samples. SPLASH should therefore be able to perform bacterial strain typing *de novo*. This is important for surveillance: tracing outbreaks and monitoring the evolution of key traits such as virulence and antibiotic resistance. Current approaches to strain typing are time-consuming and hinge on reference genomes and annotated strain-defining genetic variants. Phylogenetic models rely on SNPs to estimate the highest-likelihood trees; however, these models require manual scrutiny and are inconsistent with key mechanisms of bacterial evolution, namely extensive horizontal gene transfer (HGT).

We tested if SPLASH could perform strain typing by assessing its ability to classify lineages within the microbial species of *Mycobacterium tuberculosis*, *Listeria monocytogenes*, and Group B *Streptococcus* (GBS) (among others). These species have well-defined phylogenetic structure and possess distinct forms of genome plasticity. *M. tuberculosis* has rare HGT and a mutation rate folds lower than most bacterial species, whereas *Listeria* and GBS have high rates of HGT.^38–41^ Here we show that SPLASH performs rapid, automatic phylogenetic classification not driven by library or collection or “batch effects,” including across large-scale clinical studies and across multiple continents (Figure 5).

**Figure 5.**
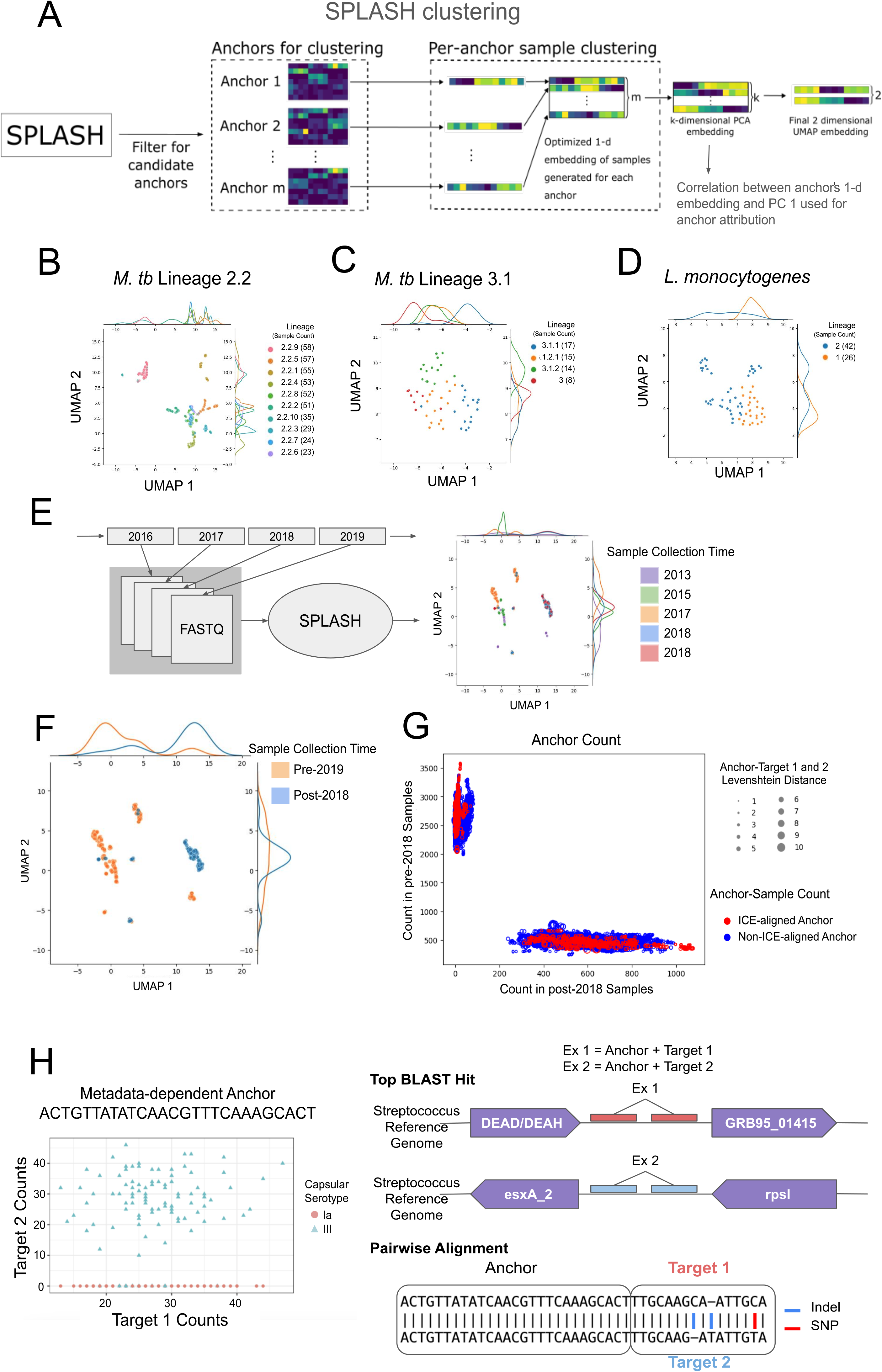
Reproducible *de novo* phylogenetic modeling and detection of strain variation. (A) For each SPLASH-called anchor, SPLASH reports a single vector, called a sample embedding, representing the optimal partitioning of samples by their target distributions. In SPLASH clustering, anchors’ sample embeddings are aggregated and PCA performed on the resulting matrix (formally, its transpose), denoising individual anchors’ sample-target distributions and building statistical strength. To represent sample similarity in 2 dimensions, UMAP is performed on the matrix resulting from PCA. Applied to microbial isolate sequencing data, metadata-blind SPLASH clustering groups samples by lineage and clade completely *de novo*. (B) *M. tb* lineage 2.2: UMAP of 20D sample embedding achieves 81% 5-NN accuracy. (C) *M. tb* lineage 3.1: UMAP of 10d sample embedding achieves 87% 5-NN accuracy. (D) *L monocytogenes*: UMAP of 20d sample embedding achieves 96% 5-NN accuracy. (F) SPLASH clustering of raw sequencing data from a *V. cholerae* longitudinal study yields 2 primary clusters having diverse sample collection years. (G) Labeling samples according to a collection year boundary (before or after 12:00 AM January 1, 2019) reveals highly separable clusters. Following attribution analysis, driving anchors were selected if exactly 1 of the most and second most-supported anchor-targets BLASTN-aligned to *Vibrio* Methods). Anchor counts are plotted with color indicating sample collection date category and whether the corresponding anchor-targets BLASTN-aligned to ICE. Anchors (points) sized by Levenshtein distance between anchor-targets 1 and 2 (Supplemental Methods). (H) SPLASH identifies an anchor in Group B *Streptococcus* where target fraction predicts capsular serotype. Anchor-targets 1 and 2’s top BLAST hits suggest a relationship with virulence factor (esxA_2).

SPLASH reports significant anchors along with a vector (a “sample embedding”) that can be interpreted as the weighted sample order that best reflects sample similarity in the metric of target distribution, much like the interpretation of principal components.^42^ We used the set of these vectors across anchors to develop a clustering procedure similar to our previously reported work.^42^ The clustering procedure starts with concatenating anchors’ reported sample embeddings to form a new matrix: each column corresponds to a sample and each row to an anchor (Methods). Dimensionality reduction is performed on this matrix via principal components analysis (PCA); formally we perform PCA on the transpose of this matrix. We retain the top k components, yielding a k-dimensional sample embedding, and optionally use UMAP (default parameters) on these k-dimensional sample embeddings for two-dimensional visualization. As always with UMAP, this may or may not reflect clusters that exist in higher dimensions (Methods, Figure 5A).^43^ All results below are generated using this algorithm, without any metadata or additional optimization, though such adjustments may further improve results.

We first evaluated whether unsupervised SPLASH clustering reflected known lineages, both numerically and graphically. For numerical evaluation, we used k=5 nearest-neighbors (k-NN) classification accuracy for the UMAP embeddings: for each sample, we generate a predicted label as the majority vote of its k nearest-neighbors, and compute the fraction of samples for which this predicted label matches the true label (Methods).

We performed SPLASH clustering on *M. tuberculosis* lineage 2.2 because of its impact on public health: sublineage 2.2.1 is the “modern” Beijing [sic] clade, rapidly evolving and associated with hypervirulence and drug resistance.^44^ We tested whether SPLASH could recover sub-sub-lineages such as 2.2.1 within isolates classified as lineage 2. SPLASH clustering yields 81% k-NN classification accuracy in classifying sub-lineages 2.2.1-2.2.10 (Figure 5B). SPLASH clustering performs significantly better than PCA followed by UMAP on the same raw anchor-target counts matrices: For lineage 2.2, prediction accuracy falls from 81% to 53% (Figure 5B, Suppl. Figure 5A, respectively), while for *M. tb* lineage 3.1 the accuracy falls from 87% to 61% (Figure 5C, Suppl. Figure 5F, respectively). Additionally, for *M. tb* lineage 3.1, using all anchors instead of SPLASH-called anchors, PCA followed by UMAP attains an accuracy of only 44% (Suppl. Figure 5E). We also tested SPLASH clustering on a subset of isolates classified as lineage 3: SPLASH clustering attains 87% k-NN classification accuracy (Figure 5C). For some *M. tuberculosis* isolates, the authors of the original study could not determine an isolate’s sub- or sub-sub-lineage. In such cases, an isolate was annotated with a less refined taxonomy: for example, an isolate not classifiable as lineage 3.1.1 or 3.1.2 is assigned to 3.1. To our surprise, in *M. tb* lineage 3, the first principal component of the matrix of sample embeddings orders the samples not only by lineage, but also by inferred phylogenetic distance and according to taxonomic rank (Suppl. Figure 5C, 5D).

We next evaluated SPLASH clustering on *Vibrio cholerae*, a bacterial species with higher genomic plasticity. *V. cholerae* population structure changes periodically, driven by phages, transposable elements, plasmids, and SNPs, and longitudinal sampling is used to monitor evolution of pathogenicity factors.^11^ The authors of a recent study found that circulating strains before and after 2018 experienced a shift in genetic cargo of mobile elements that conferred differential susceptibility to phage.^11^ Thus, the year of collection is expected to be partially predictable based on sequencing assays. SPLASH clustering discriminated samples collected before or after 1 January 2019 with 96% accuracy, with the samples forming two distinct clusters concordant with this metadata (per-year prediction gave 76% k-NN classification accuracy) (Figure 5E, 5F). Much of this predictive power comes from the first principal clustering direction (92% k-NN accuracy on pre vs post 2018 classification).

This striking result prompted us to perform attribution analysis for anchors driving the pre- vs. post-2018 sample clustering (Methods). Analysis of the 8,321 driving anchors showed anchors having the highest counts in samples collected before 1 January 2019 were disjoint from those having the highest counts in samples collected after 1 January 2019 (Methods, Figure 5G). Anchors with the highest counts in either pre-or post-1 January 2019 samples were found (via BLASTN) to have near-perfect identity (within 2-bp) to an assembly annotated as a *V. cholera*e SXT ICE element, where anchor-targets encoded protein-modifying variation including in an annotated potential type IV toxin-antitoxin system protein (Supplemental Methods). The authors in the 2021 study reported strain-defining changes to the gene cargo of a *V. cholerae* SXT ICE element: the exchange in a phage defense system from a restriction-modification system to a BREX system.^11^ SPLASH clustering infers strain-defining features of the SXT ICE element entirely *de novo*. Given that the 2021 study’s major finding was indeed a genetic discrimination of strains pre- and post-1 January 2019, this raises the intriguing possibility that SPLASH could be used to automatically detect temporal shifts in bacterial strains.^11^

We further tested whether strains could be classified in large studies of clinical isolates of other bacterial species. In *Listeria*, SPLASH clustering separated samples by lineage with a k-NN prediction accuracy of 96% (Lineages 1 and 2 were used as they were the only to have significant numbers of samples) (Figure 5D). In Group B *Streptococcus*, SPLASH clustering obtains 98-100% k-NN classification accuracy for each metadata tag–each of clade, clonal complex or multilocus sequence typing (MLST) tags describing the bacteria. We note that using the first 2 principal components of the embedding-aggregation method alone yields >98% accuracy in these metadata categories (Methods, Suppl. Figure 5G, 5H, 5I). Together, the performance of SPLASH clustering implies that SPLASH automatically captures features that define the diverse inter-strain bacterial strains analyzed here.

## SPLASH provides a direct statistical approach to define genetic features distinguishing strains

Finally, we investigated SPLASH’s ability to identify genetic features (e.g. SNPs or indels) distinguishing strains. Most strain-level variation in microbial populations is identified by first creating a reference genome from widely-used isolates, then performing pairwise sequence alignments. This process is multistep, time-consuming and lacks full statistical rigor. As an alternative, a simple, fully statistical approach using SPLASH can identify genetic changes that predict strains–or clusters that SPLASH identified–using a GLM regressing target fractions on metadata such as strain. We show the potential of this approach in Group B *Streptococcus*, using capsular serotype. Capsular serotype is an annotated metadata category, but it is also predicted *de novo* with high fidelity by SPLASH clustering (Suppl. Figure 5J). We tested whether the target distributions of SPLASH-called anchors could predict capsular serotype in *Streptococcus agalactiae*, or Group B *Streptococcus* (Methods).^28^ As an example of the insights possible with this approach, we identified an anchor distinguishing serotypes Ia from III based on target fractions alone. Anchor-target 1 aligns upstream of a WXG100 family type VII secretion target gene and anchor-target 2 aligns upstream of a virulence factor that resembles EsxA (a virulence factor of *Mycobacterium tuberculosis*). Pairwise alignment of anchor-target 1 and anchor-target 2 show 2 indels and a single SNP in targets 1 and 2 (Fig. 5H), supporting SPLASH’s rapid, statistical discovery potential.

In summary, SPLASH clustering operates blind to any reference genome and metadata and runs without parameter tuning between datasets, producing clusters that are highly concordant with known metadata. Moreover, it can assign genomic features consistent with these differences through a simple general linear model (Methods), which include sequences absent from the reference assembly.^28^ Future work will further investigate and expand SPLASH’s clustering method and study the biological significance of these unaligned sequences.

## Conclusion

SPLASH and a new statistical approach to de novo assembly–compactors–are a user-friendly and highly efficient method for performing microbial sequence analysis, broadly defined. They are computationally efficient and require no reference genome or metadata to generate a variety of predictions that today are approached with disparate and subfield-specific bioinformatics tools. For example, SPLASH and compactors automate the process of discovery of new genome diversity, while also performing more classical tasks that historically require assembly and alignment, such as strain typing.

This work shows that statistical analysis of massive quantities of raw reads (in the billions per study), and this analysis alone–without any reference genome or database–can generate new predictions with characteristics of novel MITEs and CRISPR repeats and suggest associations between domains of unknown function and transposition. Beyond this, a first application of SPLASH clustering achieves surprising precision in microbial strain typing without algorithmic tuning. Bacterial species have highly diverse mutational spectra and rates including differential occupation with mobile genetic elements, horizontal gene transfer, and rates of recombination–*M. tuberculosis* having a slow mutational clock and relatively few (if any) known examples of horizontal gene transfer, and other bacteria such as Group B *Streptococcus* having extremely high rates. The same clustering algorithm–with no parameter tuning–achieves high predictive power for strain clustering of both species and further enables attribution analysis to these differences bypassing mapping, without the requirement of a reference genome. By design, SPLASH and compactors perform these tasks multiple-fold (or more) more quickly than state-of-the art. SPLASH and compactors’ low human and compute resource requirements suggest that massive analysis of the microbial world with these tools is possible, and that they can be configured to automate discovery.

## Acknowledgments

We thank members of the Salzman lab for feedback, Matt Durant for suggestions on benchmarking for compactors and sharing simulation regimes, Andrew Fire, Silvana Konermann, Patrick Hsu and John McSpedon for useful discussions. T.Z.B. is supported in part by the Stanford Graduate Fellowship, the NSF GRFP, and the Eric and Wendy Schmidt Center at the Broad Institute of MIT and Harvard. A.F.C. is supported in part by DP2AI171122. J.S. is supported in part by 1R35GM139517-01.

## Supplemental Files

**Suppl. File 1.** (A) For each dataset, the number of SPLASH-called anchors, SPLASH peak memory usage and runtime, sum of downloaded FASTQs’ compressed size, sum of downloaded FASTQs’ read counts (line count / 4), and total number of downloaded FASTQs. (B) For each dataset, the compressed size and the uncompressed line count / 4 of each FASTQ. All FASTQs input to SPLASH for each dataset are represented.

**Suppl. File 2.** (A) BLASTN-alignment of anchors in Table 3. (B) Compactors for anchors in (A) having no BLASTN alignment with e-value < 0.05 (Methods). (C) Compactors from (B) investigated for repetitive element and (D) their BLASTN alignments to the NCBI nr/nt database. (E) SRA BLAST hits for the repetitive element in ERR6713714, ERR6713710, ERR4025905, ERR6712627. (F) Reads acquired using Pebblescout and SRA BLAST. (G) BLASTx and (H) Pfam alignment of reads acquired using Pebblescout and SRA BLAST.

**Suppl. File 3.** (A) The 10,000 highest-entropy SPLASH-called anchors in *V. cholerae* excluding anchors Bowtie2-aligning to UniVec or Illumina adapter sequences. (B) The *V. cholerae* superintegron repeat anchor’s per-sample anchor-target counts. (C) BLASTN results for those of the *V. cholerae* superintegron repeat anchor’s anchor-targets having > 0.1% of the total counts. (D) Multiple-sequence alignment of the extracted inter-repeat regions adjacent to the *V. cholerae* superintegron repeat anchor’s target 1. (E) Multiple sequence alignment in (D) cropped to produce a consensus with occupancy of at least 80%. (F) Multiple sequence alignment of sequences in (D) *in silico* translated in frame 3 to the NCBI database proteins generating the best 4 BLASTx hits to the consensus in (D). (G) GenBank flat file containing gene predictions for the repeat array in CP077425.1. (H) GenBank flat file containing gene predictions for the repeat array in CP017798.1. (I) Compactors and all Pfam alignments having e-value < 0.05 for the *V. cholerae* anchor and its reverse-complement.

**Suppl. File 4.** (A) Compactors for analysis of select anchors in tables 3 and 4. (B) Pfam hits of compactors in (A). (C) BLASTN results of Pfam-aligned compactors in (A). (D) 80th percentile of entropy and effect size for Pfam domains having >= 10 upstream compactors unique at a given extension iteration. (E) Compactors for anchors selected for analysis, regenerated to achieve greatest sensitivity.

**Suppl. File 5.** (A) FASTA containing an anchor’s upstream compactor Pfam-aligned to LRR_5. (B) Result of BLASTx search of the compactor in (A). Compactors were generated for an LRR_5-associated anchor and its reverse-complement in single samples (Methods). (C) Compactor-sample counts matrix for the anchor in (A) for compactors generated from the anchor’s reverse-complement and having at least 5 counts. (D) The best BLASTN hit for each compactor in (C). (E) Compactor-sample counts matrix for the anchor in (A), for compactors generated from the anchor and having at least 5 counts. (F) The best BLASTN hit for each compactor in (E). (G) Anchor-target-sample counts matrix for the anchor in (A), for anchor’s reverse-complement’s anchor-targets having at least 5 counts. (H) The best BLASTN hit for each anchor-target in (G). (I) Anchor-target-sample counts matrix for the anchor in (A), for anchor’s reverse-complement’s anchor-targets having at least 5 counts. (J) The best BLASTN hit for each anchor-target in (I).

## Methods

### Datasets included in the study

SPLASH was run on R1 FASTQs described below and listed in Supplemental File 1.

#### Capsule Intestinal Profiling (Shalon et al. 2023)

302 Illumina NovaSeq 6000 FASTQs (PRJNA822660) sampling stool, saliva, and intestinal fluid, the latter using dissolvable devices ingested by the 15 study participants (Suppl. File 1).

#### Streptococcus pneumoniae (Chaguze et al. 2020)

909 Illumina HiSeq 2000 FASTQs (PRJEB3084) sampling the central nervous system (CNS), cerebrospinal fluid, and non-CNS tissues including blood, joint fluid, and knee, pleural, and lung aspirate of 98 infants in 11 African countries (Suppl. File 1).

#### Group B Streptococcus (Chaguza et al. 2022)

1,512 Illumina HiSeq 2000 GBS isolates (PRJEB14124) collected in a 30-year longitudinal study of infants in the Netherlands with acute invasive disease (Suppl. File 1).

#### Group A Streptococcus (Kachroo et al. 2019)

2,101 Illumina NextSeq 500 GAS isolates (PRJNA434389) collected in a longitudinal study of invasive infection in Canada, Denmark, Finland, Iceland, Norway, and the United States from 1991 through 2016 (Suppl. File 1).

#### Group A Streptococcus (Southon et al. 2020)

1,515 Illumina NextSeq 500 GAS macrolide-resistant clinical isolates (PRJNA614628) collected in Iceland from 1998 to 2016 (Suppl. File 1).

#### Listeria (Maury et al. 2016)

69 Illumina HiSeq 2000 FASTQs (PRJEB10802) sequencing gDNA of 69 *L. monocytogenes* strains, sampled from human clinical infection (37), animal infection (16), food (12), environment (1) and unknown origin (3) (Suppl. File 1).

#### E. coli (GenomeTrakr, PRJNA230969)

270 Illumina MiSeq FASTQs sampling *E. coli*, produced by the GenomeTrakr network (Suppl. File 1).

#### V. cholerae (LeGault et al. 2021)

283 Illumina HiSeq 4000 WGS FASTQs (PRJNA723557) sampling *V. cholerae* clinical isolates in a study of SXT ICE-mediated phage resistance (Suppl. File 1).

#### Inflammatory Bowel Disease Multi-omics Database (IBDMDB)

1,338 FASTQs (PRJNA398089) of Illumina HiSeq 2000 or 2500 metagenomic sampling in a longitudinal study of 50 Crohn’s disease patients and 30 ulcerative colitis patients (Suppl. File 1).

#### Comprehensive Resistance Prediction for Tuberculosis: an International Consortium (CRyPTIC)

SPLASH was run separately on 450 (lineage 2) and 54 (lineage 3) FASTQs of *M. tuberculosis isolates*, sequenced using Illumina hardware (Suppl. File 1). Data were originally generated in a study of drug resistance in 6,814 *M. tuberculosis* isolates.

### SPLASH runs

For all datasets, SPLASH version 2.0.3 was run with the following command:

~~~
splash --outname_prefix result --anchor_len 27 --gap_len 0 --target_len 15
       --poly_ACGT_len 8 --dump_Cjs --max_pval_opt_for_Cjs 0.05
       --with_effect_size_cts --with_pval_asymp_opt --n_threads_stage_1 8
       --n_threads_stage_2 32 --n_bins 32 --n_most_freq_targets 10
       --kmc_use_RAM_only_mode --kmc_max_mem_GB 6
       --dump_sample_anchor_target_count_txt
       --dump_sample_anchor_target_count_binary.
~~~

### Tables 1a-k, Bowtie2 alignment of high effect size and highest-entropy anchors

#### Significant anchors selected by effect size and target entropy

SPLASH reported at least 1 million significant anchors in 9/10 datasets, so anchors were subsampled by the Shannon entropy of their target count distributions (*target_entropy*) and the degree to which these distributions in samples could be partitioned into 2 sets (*effect_size_bin*).^42^ Anchors in each dataset were selected (from results.after_correction.scores.tsv) if having *effect_size_bin* >= 0.9 and *target_entropy* >= 0.9, then were ordered by descending *target_entropy* and the top 10,000 (or all, if fewer than 10,000 remained after filtering) taken into Table 1. SPLASH statistics were added for interpretability.

#### Filtering contaminant sequences

Bowtie2 v2.5.0 was used to align anchors to UniVec and Illumina adapter sequences (Supplemental Methods).^45^ Anchors mapping to either reference were discarded, as were those for which either of the top 2 targets mapped to UniVec or Illumina adapter sequences. Anchors were also removed if they or either of their top 2 targets reported homopolymers of length > 7.

#### Bowtie2 annotation of anchors

Anchors and their targets in Table 1 were mapped to reference databases (Supplemental Methods) using the following command (Bowtie2 v2.5.0):

~~~
bowtie2 -f -p 32 --quiet --reorder -x <INDEX> -U <anchors.fasta>.
~~~

Bowtie2-alignment summary files were generated, reporting for each assembly’s subheader: the name of the assembly, the subheader, the number of anchors aligning to this subheader, and the number of anchors aligning only to this subheader. To summarize anchors’ Bowtie2 alignments, the *weighted_aligned_targets_value* was computed; this is defined as the sum of an anchor’s aligned targets’ read support divided by the total anchor read support.

### Kolmogorov-Smirnov test

To perform the Kolmogorov-Smirnov test, we used scipy.stats.ks_2samp, SciPy v1.9.1.^46^

### glmnet

To identify anchors having phenotypic relevance, we ran glmnet multinomial regression using provided metadata.^28^ Anchors were filtered to select those appearing in more than 5% of samples and having *avg_hamming_distance_max_target* > 5. Selected anchors were processed if in the top 100,000 for a given dataset by *effect_size_bin*.

### Algorithm for compactors generation

The algorithm takes as an input a set of anchors of length s (e.g., provided by SPLASH). For every occurrence of an anchor in a sample set, a sequence of N consecutive non-overlapping *k*-mers (encoded as 64-bit integer numbers) is returned, with N and k being the algorithm parameters (Suppl. Figure 6A). If a read is shorter, the maximum possible number of *k*-mers is output. Then, the *k*-mers at positions 0, k, 2k, …, (N-1)k following the anchor are investigated independently to identify “trusted” *k*-mers, i.e., those representing real sequence diversity (Suppl. Figure 6B). Starting from the most abundant *k*-mer (which is always considered as trusted) the algorithm greedily identifies other trusted *k*-mers that were unlikely to be produced as a result of sequencing error. The procedure uses a probabilistic model considering the *k*-mer abundance together with the Hamming distance d to the currently investigated trusted *k*-mer. In particular, the algorithm upper bounds the probability of a candidate *k*-mer to be produced by the sequencing error ε using the Poisson distribution:

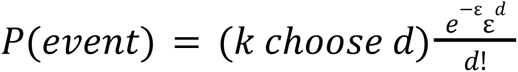

The decision rule to recognize the candidate as a result of sequencing error takes into account its abundance, the abundance of the currently analyzed trusted *k*-mer and scaling factor β:

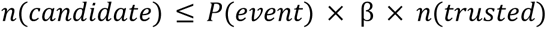

If an already existing trusted *k*-mer satisfies the condition above, the candidate is considered as a result of a sequencing error produced from this trusted *k*-mer. Consequently, if none of the trusted *k*-mers meets the condition, the candidate represents a real sequence and is added to the set of trusted *k*-mers. An additional parameter *lower_bound* indicates how many times a *k*-mer must appear in the reads at the currently considered distance from the anchor to be included as a “trusted” *k*-mers. Therefore, sequence fragments with a support in reads below *lower_bound* will not generate a compactor.

Once trusted *k*-mers are established, each *k*-mer can be then assigned with its representative, i.e., the closest in terms of the Hamming distance trusted *k*-mer (trusted *k*-mer is its own representative). Additionally, each trusted *k*-mer can be assigned with its support, i.e. the number of its exact occurrences (*exact_support*) summed with the number of non-trusted *k*-mers it represents.

After determining trusted *k*-mers, the tuples of N consecutive *k*-mers following the anchor are analyzed. In particular, non-trusted *k*-mers are replaced with their representatives and for each tuple, the sum of corresponding Hamming distances is recorded. Tuples with the sum lower than *max_mismatch* (algorithm parameter) are compactor candidates. The candidates are then sorted with respect to the number of occurrences and final compactors are identified as “trusted tuples,” by a probabilistic procedure similar to the one used for identifying trusted *k*-mers (Suppl. Figure 6C). Note, that due to replacing *k*-mers with their representatives, compactors do not necessarily appear in the reads exactly (their exact support can equal to 0) unlike the *k*-mers composing them.

Finally, the obtained compactors are subject to recursive extension. Namely, the last s-mer of a compactor (referred to as an extender) is used as an anchor and the aforementioned steps are repeated (Suppl. Figure 6D). The extension is performed only on compactors whose extenders are sufficiently specific for the anchor they originate from (i.e., the original anchor has to appear in at least *min_extender_specificity* fraction of s-mers preceding the extender in the reads at the anchor position).

### Compactors assessment: long assembly reconstruction

ART (v2.5.8) was used to simulate Illumina reads of different lengths (75, 100, 150, 200, 250) with various sequencing coverage (2, 5, 10, 20, 40, 60, 50, 100, 150) from a 25-kbp random reference.^21^ The built-in error models for HiSeq (read length up to 150-bp) and MiSeq (read length greater than 150-bp) devices were used. The command line was as follows:

~~~
art_illumina -ss {device} -i {reference FASTA} -l {read length} -f {cov} -o {output FASTQ}
~~~

with *device* being HSXt (HiSeq) or MSv1 (MiSeq).

Then, an attempt was made to reconstruct the middle 5-kbp of the reference (10,000:15,000) by generating compactors starting from a 27-base anchor at position 10,000 in the genome. Compactors (v2.3.0) were tested for *min_extender_specificity* varying from 0.5 to 0.9 using the following command:

~~~
compactors --kmer_len 27 --num_kmers 1 --lower_bound 1 --max_length 5000
           --epsilon 0.001 --beta 0.5 --min_extender_specificity
           {min_extender_specificity value} {FASTQ list} {one element anchor list}
           {output}.
~~~

Compactors were compared with SSAKE (v4.0) and MEGAHIT (v1.2.9). ^22,23^ As SSAKE was not able to reconstruct the assembly when using the anchor as a seed, a default mode which generates seeds from a read set was employed:

~~~
SSAKE -p 0 -f {empty FASTQ} -g {FASTQ file} -i 1 -w 1 -m 20 -o 1 -c 1 -r 0.51 -b
{output}.
~~~

MEGAHIT was run with the following command:

~~~
megahit -r {FASTQ file} -o {output} --min-contig-len 27 --min-count 2.
~~~

Simulations were repeated 30 times for every experimental condition (read length, sequencing coverage, algorithm configuration).

### Compactors assessment: precision-sensitivity trade off

Ten *Vibrio cholerae* samples (SRR14297458-67) were used to establish the amount of false assemblies (Suppl. Figure 7). As a first step we identified 27-base anchors from the reads using SPLASH and selected 500,000 with the largest effect size. Then, every read in a set was split in half. Resulting set of half-reads was provided as an input to compactors (v2.3.0) with the following parameters: k = 27, N = 1, and maximum compactor length of 108-bp, which corresponds to 2 extensions. Finally, we verified the presence of compactors in the original, full-length reads to establish a number of true positive (TP) and false positive (FP) discoveries. True positive discoveries were defined as sequences represented more than once in the full length reads. The TP/FP trade off was controlled by altering *lower_bound* parameter from 2 to 10:

~~~
compactors --kmer_len 27 --num_kmers 1 --max_length 108
--lower_bound {lower_bound value} --num_threads 16
{half-reads FASTQ list} {anchors list} {output}.
~~~

Compactors were compared to SSAKE (v4.0) and MEGAHIT (v1.2.9) by running the algorithms on the split reads and searching the resulting contigs for the fragments of length 108-bp starting with an anchor. These fragments were then identified in the ground truth to establish TP and FP counts. In the case of SSAKE, the anchors were given as seeds and minimum base ratio parameter (-r) was altered from 0.51 to 0.99 to control FP/TP trade off:

~~~
SSAKE -p 0 -f {empty FASTQ} -g {half-reads FASTQ} -s {anchors FASTA} -i 0 -w 5 -m 20
      -o 1 -c 1 -r {r value} -b {output}.
~~~

Controlling precision and sensitivity of MEGAHIT was more problematic. The algorithm was run with the following command:

~~~
megahit -r {half-reads FASTQ} -t 16 -o {output} --no-mercy --k-min 29
        --k-step {k-step value} --min-contig-len 27 --min-count {min-count value}
~~~

with *k-step* = 14 and *min-count* = 1 in the most sensitive configuration and *k-step* = 12 and

*min-count* = 2 in the most precise one.

### Assessment of protein homology

#### Pfam alignment

To assess compactors’ protein homology, compactors were translated in all six frames with the standard translation table using seqkit prior to using hmmsearch from the HMMer3 package to report homologous amino acid sequences in the Pfam35 profile Hidden Markov Model (pHMM) database.^27,47,48^

#### BLASTx search

BLASTx was used with default parameters.^49^

### Compactors in Pfam domain analysis

#### Compactor generation and Pfam annotation

Anchors appearing in at least two datasets in Table 3 and/or Table 4, and all anchors from IBDMBD and the intestinal profiling study (Shalon et al. 2023) in Tables 3 and 4 were selected alongside their reverse-complements. For each dataset and its corresponding anchors and anchors’ reverse-complements, compactors were generated with the following command (SPLASH v2.1.14):

~~~
compactors --beta 0.5 --epsilon 0.001 --num_threads 20 {FASTQ list} {input anchors}
{output name}.
~~~

Compactor outputs from each dataset were joined to form Supplemental File 4A, with compactors reported if having exact_support > 0. Additional fields were added to facilitate analysis (Supplemental Methods). Compactors were submitted to Pfam search and hits retained if having a full-sequence e-value < 0.05 (Suppl. File 4B). Pfam domains were categorized based on their human-readable names (Supplemental Methods).

#### Identification of Compactors Associated With CRISPR and Mobile Elements

Compactors were selected if generated from an anchor’s reverse-complement and if unique at a given extension, thus cases where only one compactor was generated upstream of the anchor at a certain extension iteration. Pfam domains to which >= 10 such compactors aligned were selected; for such domains the 80th percentile of entropy and effect size of aligned compactors’ anchors was computed.

#### High-Sensitivity Compactors for Selected Anchors

Anchors selected for analysis were resubmitted to compactor generation to achieve the greatest sensitivity, bypassing the deduplication step in the procedure’s optimization framework. Compactors are reported in Supplemental File 4E and were generated using the following command (SPLASH v2.3.0):

~~~
compactors --beta 0.5 --epsilon 0.001 --num_threads 20 --all_anchors {FASTQ list}
{input anchors} {output name}
~~~

#### BLAST of Compactors

Pfam-aligned compactors in Supplemental File 4A were submitted to remote BLASTN search using the following command^50^:

*blastn -remote -db nt -evalue 10 -task blastn -dust no -word_size 24 -reward 1 -penalty -3 -max_target_seqs 50*.

BLASTN results were reported along with the compactors (Suppl. File 4C).

### High-Sensitivity Compactors

High-sensitivity compactors were further used in analysis of the *V. cholerae* superintegron repeat and of the highest-entropy BLASTN-unaligned anchors, where the proposed novel CRISPR repeat was identified (Suppl. Files 2B, 2C, 3I). To generate these compactors, this command was used (SPLASH v2.3.0):

~~~
compactors --beta 0.5 --epsilon 0.001 --num_threads 20 --all_anchors {FASTQ list}
{input anchors} {output name}
~~~

### Compactors Generated for Individual FASTQs

Compactor generation (SPLASH v2.1.14) was performed with beta of 0.5 and epsilon of 0.001 for an anchor and its reverse-complement to support analysis of the LRR_5-associated MITE. Compactor generation was performed for FASTQs separately; compactors having *exact_support* = 0 were removed.

### Processing offset anchors

SPLASH can call anchors that are slight offsets of each other, which can confound post-facto analyses. We apply a simple method to organize offset anchors: group together any 2 anchors that are a shift distance of at most d from each other, where two length k anchors are shift distance d from each other if the last k-d base-pairs of one anchor are equal to the first k-d base-pairs of the second anchor (and that this does not hold for d-1). This is implemented by generating a similarity matrix between the n anchors, where entry i,j is equal to 1 if anchors i and j are within a shift distance of k from each other. We identify the connected components of the graph defined by this adjacency matrix using the networkx package.^51^ To accelerate this procedure, we compare hashes of the strings to use vectorized matrix operations.

### Clustering

The clustering methods implemented here build from those devised in OASIS.^42^ To start, these methods select a subset of anchors to perform clustering with: those with significant Benjamini-Yekutieli corrected p-values (<.05), many counts (>1,000), no long runs of a single nucleotide in the anchor or dominant target (>=8), appearing in many samples (>5%), and high effect size (>50th percentile of remaining anchors after these filtering steps, max 10,000 anchors).

We utilize k-NN classification accuracy as a metric of metadata concordance: this is computed by, for each sample, examining its k-nearest neighbors in embedding space, and evaluating whether the majority of these neighbors have the same ground truth metadata label as the sample in question (performed using the scitkit-learn package).^52^ We then compute the fraction of samples for which this is true. By default, we use k=5. To assess the significance of this, we can also permute the labels of the points and compute the k-NN classification accuracy under this null model. This yields a permutation p-value, or a null mean and variance, where our predicted embedding x,y coordinates are independent of the ground truth metadata.

To cluster the samples, we generate an overall sample embedding matrix by aggregating all per-anchor sample embeddings.^42^ Concretely, SPLASH generates per-anchor sample embeddings in order to compute a p-value bound for that anchor. Embedding-aggregation takes these per-anchor sample embeddings and stacks them in a matrix to generate an anchor-by-sample embedding matrix. Then, to obtain low-dimensional embeddings and aggregate information across anchors, PCA is applied to this matrix (formally, the transpose of this matrix). We refer to the first principal component (principal right singular vector of the untransposed matrix) as the first embedding direction identified by SPLASH. In some plots this is left unnormalized (not rescaled to be a unit vector). As shown in Figure 5A, a k-dimensional embedding is generated for the samples, which is optionally reduced to a 2-dimensional embedding using UMAP, for visualization purposes. Automated methods can be devised to select k. In this work we utilize k=2 (no UMAP) for easily separable datasets, and choose the better performing clustering between k=10 and 20 (k=10 works well, but k=20 yields improved performance for datasets with more complex cluster structure such as *M. tuberculosis* lineage 2). UMAP was run with default parameters (number of nearest neighbors=15), and empirically yielded clusterings that respected the clustering structure of the data better than PCA (just applying embedding-aggregation).

#### Attribution analysis

In order to identify anchors that are concordant with the different clustering directions identified by PCA, we utilize a form of attribution analysis. This works by recording, for each anchor used for clustering, the inner product between its sample embedding and the top k embedding-aggregation vectors (k=5 by default), the principal components. If an anchor has a high absolute inner product with an output embedding-aggregation vector, then we say that it is “driving” the clustering in this direction. In *V. cholerae* analysis, driving anchors were defined as those having absolute inner product with the first principal component > 0.9.

To test the performance of naive methods, we look at the raw counts tables for the anchors used for clustering, and run a simple clustering algorithm on these. We generate an extended contingency table by stacking the contingency tables of each anchor.^42^

## Data and code availability

All FASTQ records were downloaded from the NCBI SRA. SPLASH software used in this study is freely available at: https://github.com/orgs/refresh-bio/packages/container/package/splash.

---

**Suppl. Figure 1.**
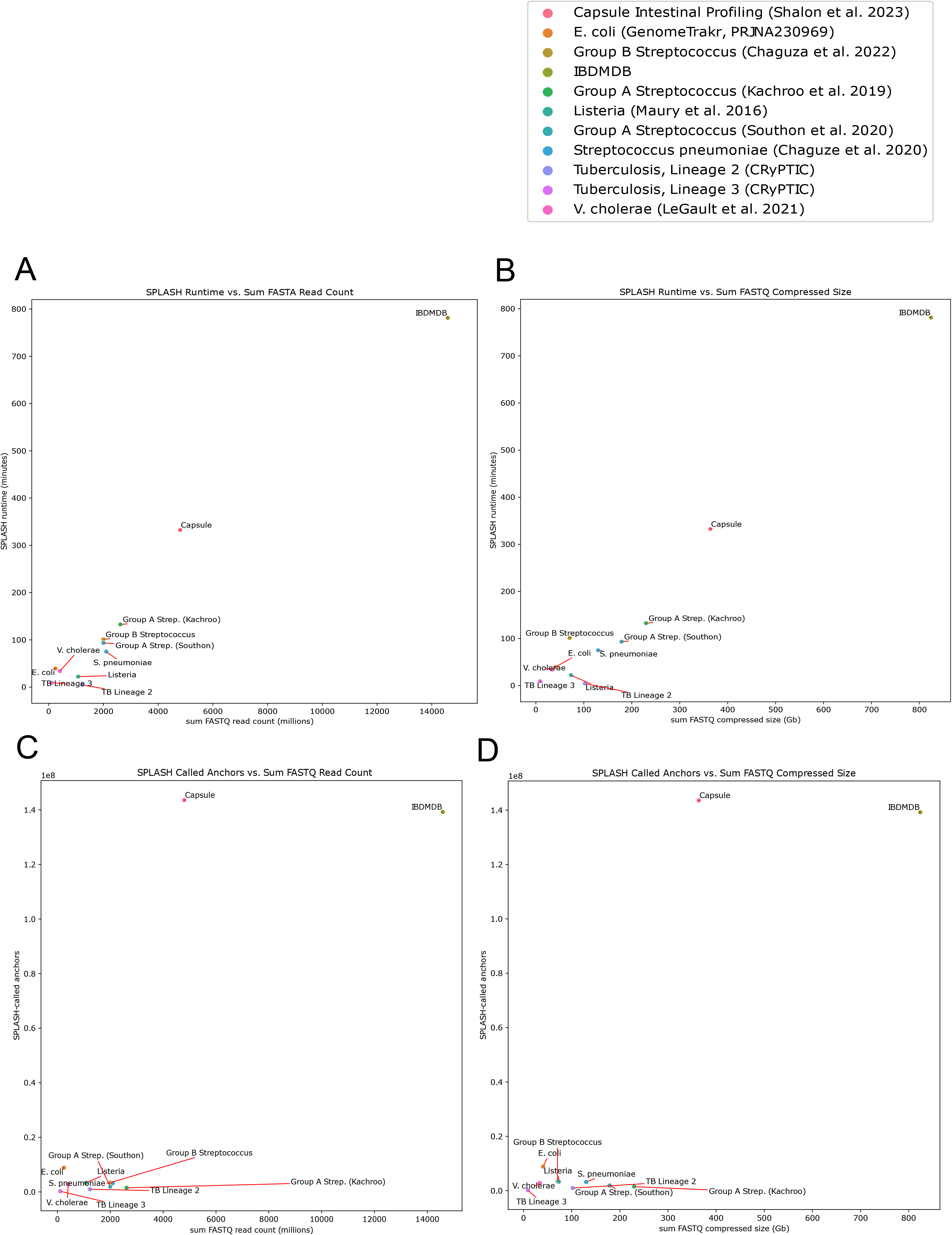
(A) SPLASH runtime (minutes) and the sum FASTQ read count (millions) for each dataset. (B) SPLASH runtime (minutes) and the sum FASTQ compressed size (Gb) for each dataset. (C) The number of SPLASH-called anchors (hundreds of millions) and the sum FASTQ read count (millions) for each dataset. (D) The number of SPLASH-called anchors (hundreds of millions) and the sum FASTQ compressed size (Gb) for each dataset.

**Suppl. Figure 2.**
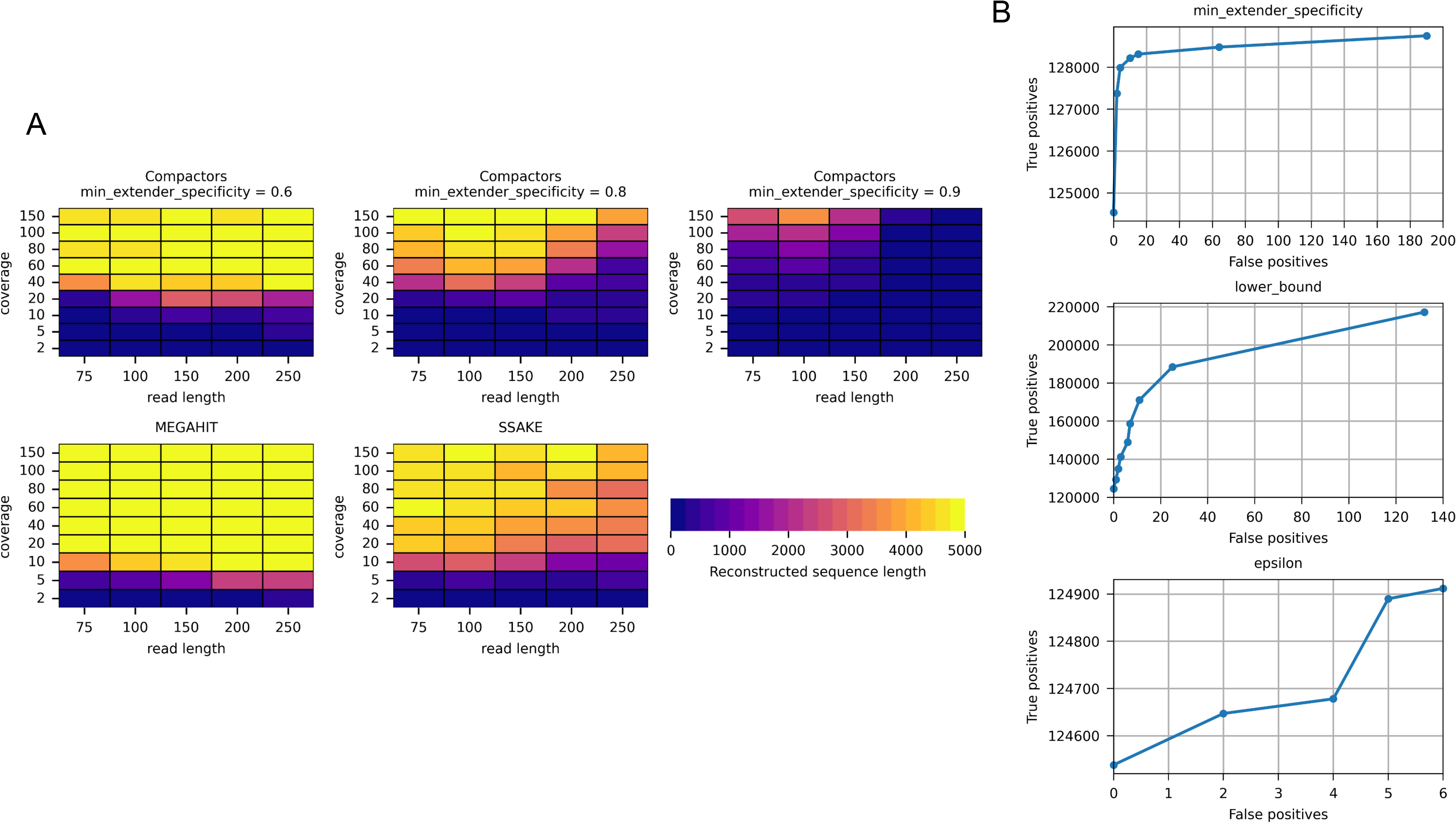
Performance of compactors on synthetic and real data. (A) The reconstruction of a middle 5-kbp sequence from a random 25-kbp reference using ART-simulated Illumina reads of different lengths. MEGAHIT and SSAKE assemblers were included in the comparison. (B) The effect of compactor generation parameters on the true and false positive trade-off. Compactors generated from *Vibrio cholerae* reads split in half were checked for presence in the full-length reads (unique sequences were removed from the ground truth as possible sequencing artifacts).

**Suppl. Figure 3.**
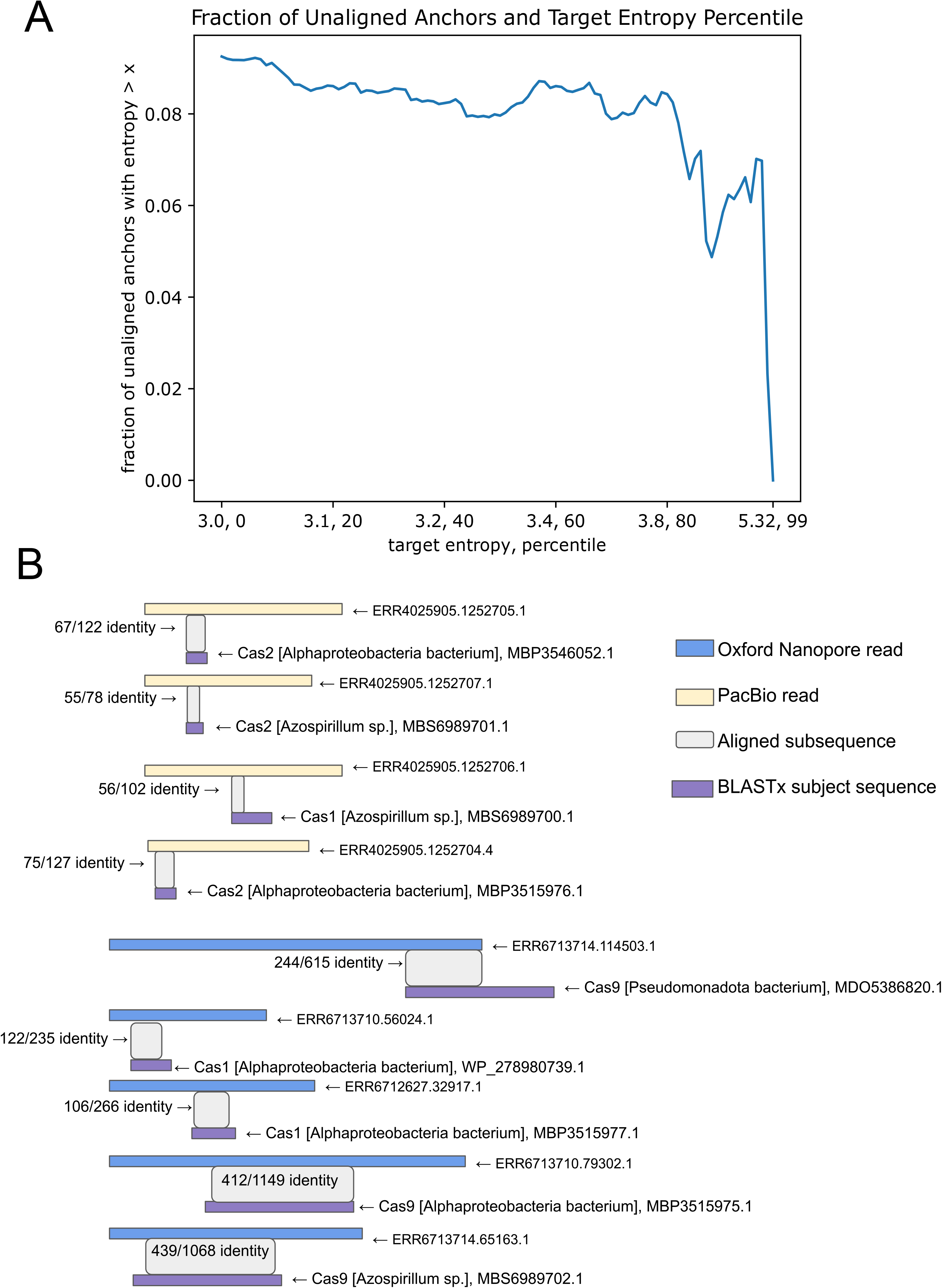
(A) Anchors in Table 3 defined as unaligned if lacking BLASTN alignment to NCBI nr databases with e-value < 0.05 (Methods). For each percentile of entropy x in [0, 99], the fraction of unaligned anchors in percentiles > x with respect to the total unaligned anchor count. (B) For the 9 reads long reads reporting the proposed novel CRISPR repeat, the best BLASTx alignment to a Cas protein is visualized.

**Suppl. Figure 4.**
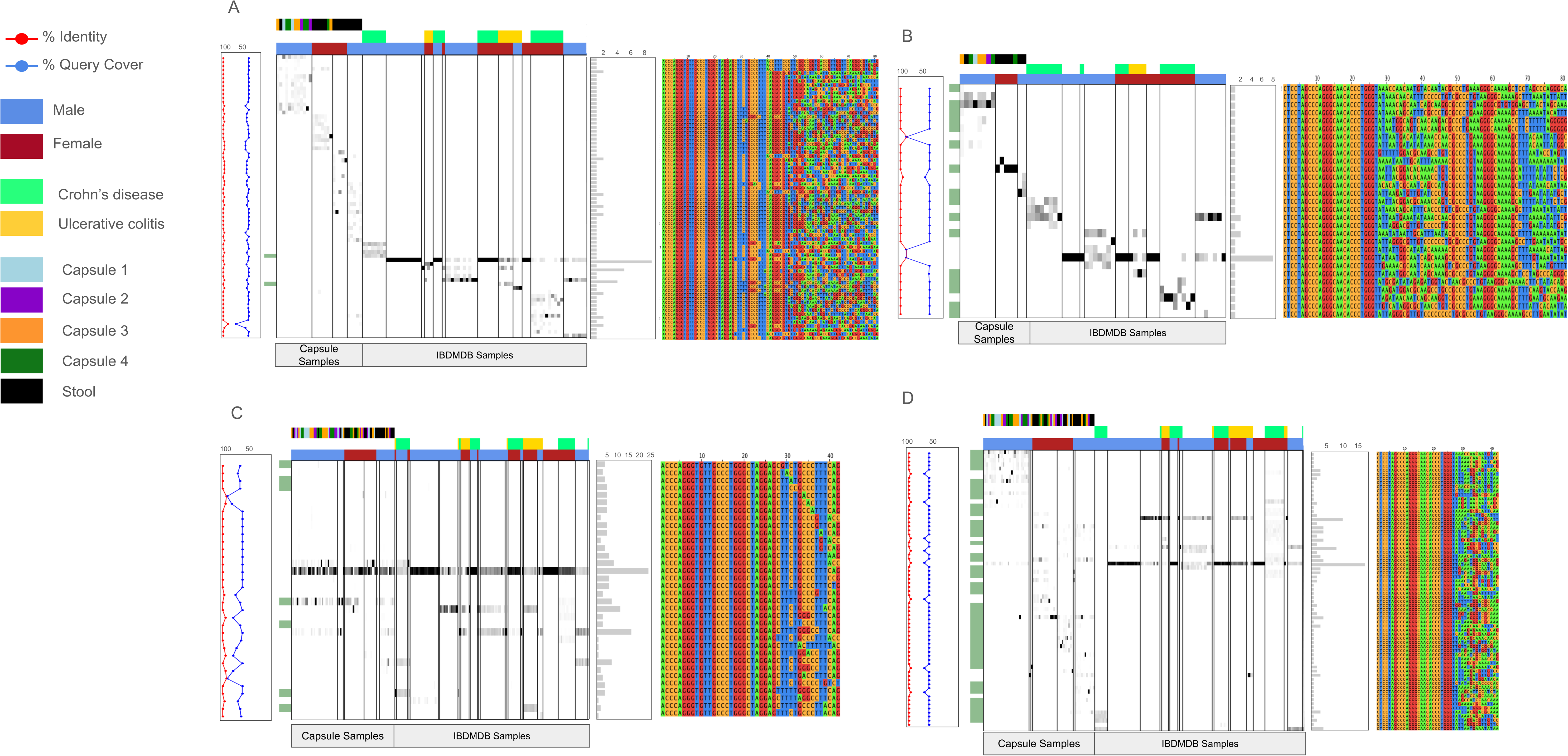
Heatmaps of anchor-target or compactor sample-count fraction with vertical bars separating donor. The top-marginal color-bars indicating sex, disease, and collection location metadata. If only the anchor, and not the compactor, is BLASTN-aligned, the left-marginal color-bars show green. A connected dot-plot reports BLASTN % query cover and identity in blue and red, respectively. At right, a multiple sequence alignment and barplot reports the number of distinct donors reporting the sequence. For an anchor having an upstream compactor aligned to LRR_5: (A) downstream and (B) upstream compactors and (C) downstream and (D) upstream anchor-target sample-count fractions and additional data are presented as described.

**Suppl. Figure 5.**
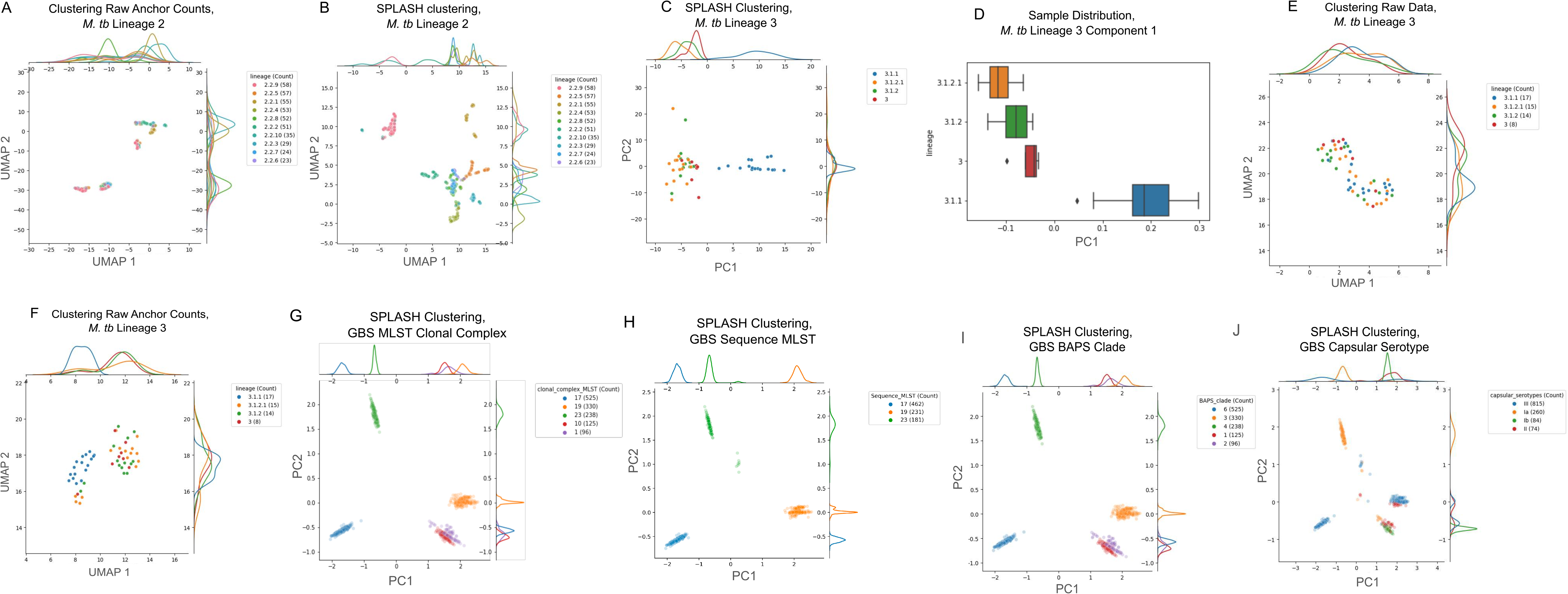
(A) PCA and UMAP-clustering of raw anchor-target counts matrices for significant anchors from SPLASH yields 53% k-NN classification accuracy in *M. tuberculosis* lineage 2. Performed for PCA embedding dimensions k of [10, 20, 50, 100, 200] input to UMAP, the maximum accuracy was attained at k=100. (B) In *M. tuberculosis* lineage 2, SPLASH clustering for PCA dimension k=20 yields an embedding with much higher k-NN classification accuracy than in Suppl. Figure 5A, at 81%. k=10 principal components are insufficient for expressing the complex clustering structure, yielding only 64% k-NN classification accuracy (not shown). (C) SPLASH clustering of *M. tuberculosis* lineage 3 yields 63% k-NN accuracy simply using principal components 1 and 2. (D) Univariate distributions of samples along principal component 1 (from Suppl. Figure 5C) illustrate that the embedding stratifies samples by lineage classification and by certainty of classification. (E) Using anchor-target counts for all anchors, not just those called by SPLASH, a standard clustering workflow of PCA followed by UMAP yields poor clustering performance, with a k-NN accuracy of 44% (maximum accuracy was attained after testing with [10, 20, 50, 54] PCA embedding dimensions input to UMAP). (F) PCA and UMAP of raw counts matrices for SPLASH-called anchors. As with (E) and (A) this simplified method falls short, with 61% k-NN accuracy (max accuracy taken over [10, 20, 50, 54] PCA embedding dimensions). (G-J) Clustering with principal components 1 and 2 in Group B *Streptococcus* yields k-NN prediction accuracy of (G) 98% in MLST clonal complex, (H) 100% in sequence MLST, (I) 98% in BAPS clade, and (J) 96% in capsular serotype.

**Suppl. Figure 6.**
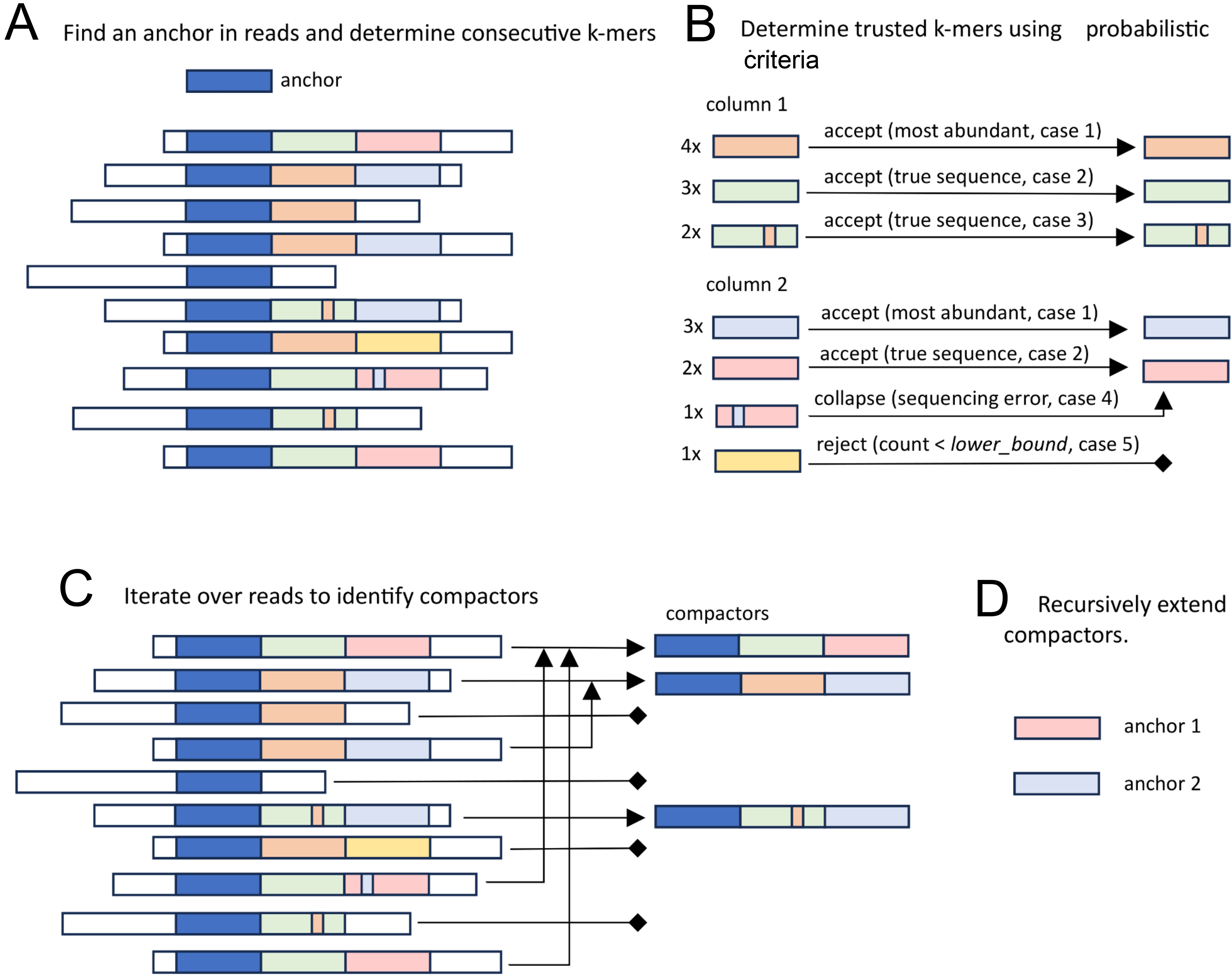
Generation of compactors for a given anchor. (A) Finding the anchor in the reads and determining following *k*-mers. Sequence fragments of different colors are highly dissimilar in terms of the Hamming distance. (B) Applying probabilistic model to identify *k*-mers present in the reads (referred to as trusted *k*-mers) with several cases being shown. Case 1: most abundant *k*-mer is always considered trusted. Case 2: Abundant *k*-mer highly dissimilar to the previous true *k*-mers considered by the model as trusted. Case 3: *K*-mer similar to one of the previous trusted *k*-mers sufficiently abundant to be considered trusted. Case 4: Low abundant *k*-mer similar to one of the previous trusted *k*-mers considered as a sequencing error and collapsed with its predecessor. Case 5: *K*-mer highly dissimilar to the previous trusted *k*-mers occurring less then lower_bound times is rejected. (C) Identifying compactors by iterating over reads and taking sequences of trusted *k*-mers representing them. Reads assigned with the same sequence of trusted *k*-mers share a compactor. (D) Recursive extension of compactors by using their ending *k*-mers as new anchors.

**Suppl. Figure 7.**
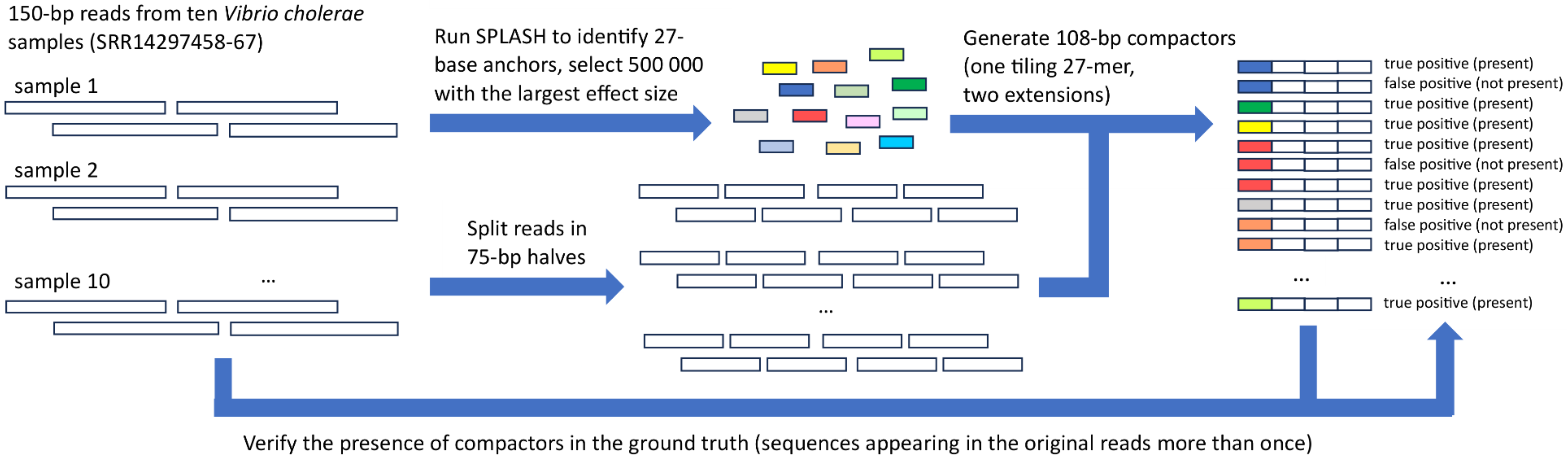
The experimental pipeline for quantifying false positive assembly artifacts produced by compactors.

