## Supplemental Methods for "Ultra-efficient, unified discovery from microbial sequencing with SPLASH and precise statistical assembly"

### Bowtie2 annotation of anchors

Bowtie2 indices used for alignments reported in Table 1 were generated using the following FASTA records, where hits to these records use the given name: *aedes*: GCF_002204515.2_AeagL5.0; *dfam_te_eukaryota*: Dfam v3.5[1] ; *direct_repeats* and *spacers*: resp. CRISPRCasdb direct repeats and spacers databases,[2] downloaded in October 2021; *ebola*: GCA_900094155.1_ASM90009415v1; *ECOLI_CFT073*: GCF_000007445.1_ASM744v1; *e_coli* and *ECOLI_K12*: GCF_000005845.2_ASM584v2; *escherichia_phage_phiX174*: NC_001422.1; *eukaryota_its1_itstonedb*: ITSoneDB,[3] downloaded in December 2021; *eukaryota_its2_biozentrum*: Internal Transcribed Spacer 2 Ribosomal RNA Database (ITS2),[4] downloaded in December 2021; *grass_carp*: GCF_019924925.1; *GrCh38p14_GCF_000001405*: Human reference genome GRCh38.p14, GCA_000001405.40; *HIV_GCF_000856385*: GCA_000856385.1; *ice_iceberg*: ICEberg 2.0,[5] downloaded in December 2021 (https://bioinfo-mml.sjtu.edu.cn/ICEberg2/download/ICE_seq_all.fas); *illumina_adapters*: sequencing adapters commonly used by Illumina, downloaded from TrimGalore[6] in October 2021; *InfluenzaA*: GCA_001014065.1; *listeria*: GCA_000196035.1_ASM19603v1; *MERS*: GCF_000901155.1_ViralProj183710; *mge_aclame_genes_all_0*: mobile genetic elements from ACLAME,[7] downloaded in December 2021; *MonkeyPox*: GCA_023913615.1; *narnavirus*: Bowtie2 index generated using a FASTA containing contigs MW434196.1 and MW434168.1; *octopus*: GCF_001194135.2_ASM119413v2; *octopus_trinity*: O. bimaculoides assembled transcriptome[8]; *RF_all*: Rfam[9] FASTA records downloaded on 3 July 2021 via wget ftp://ftp.ebi.ac.uk:21/pub/databases/Rfam/CURRENT/fasta_files/*; *sars*: GCF_009858895.2; *StrepA*: GCA_900475035.1_41965_F01; *StrepB*: GCA_003966545.1_ASM396654v1; *StrepP*: GCA_000019005.1_ASM1900v1; *TAIR10*: The Arabidopsis Information Resource (TAIR)[10] TAIR10 genome assembly; *TB_h37rv*: NC_000962.3; *tncentral_te_prokaryotes_final*: prokaryotic transposons from the TnCentral database,[11] downloaded in December 2021; *UniVec*: vector contaminant sequences, downloaded in November 2021[12]; *vibrio_cholerae*: GCA_012275105.1_ASM1227510v1 (used to annotate anchors including that matching the *V. cholerae* superintegron repeat); *WBcel235*: Caenorhabditis elegans reference genome GCA_000002985.6; *wolbachia*: Wolbachia pipientis wAlbB genome GCA_004171285.1_ASM417128v1; *zostera_marina*: GCA_001185155.1_Zosma_marina.v.2.1.

### Pfam domain analysis

*Added fields to facilitate analysis of compactors up- and downstream of seeding anchors*

*is_RC* indicates whether an anchor is the reverse-complement of a SPLASH-called anchor in the corresponding dataset; *RC_called_also* indicates whether an anchor was called both as itself and its reverse-complement in the corresponding dataset; *compactor_count* is the number of distinct compactors for a given combination of anchor, dataset, and *num_extended*. *max_compactor_count* is the maximum value of compactor_count for a given anchor in a given dataset. *rc_compactor_count* and *rc_max_compactor_count* are the corresponding values for an anchor's reverse complement in a given dataset: for a given anchor called in IBDMDB at *num_extended* 1, *rc_compactor_count* for this anchor at *num_extended* 1 in IBDMDB equals the anchor's reverse-complement's *compactor_count* at *num_extended* 1 in IBDMDB, and *rc_max_compactor_count* will be the *max_compactor_count* for the anchor's reverse-complement in IBDMDB.

*Human-readable Pfam domain names*

Pfam domains were categorized using human-readable domain names: transposase if containing Tnp or transposase (case-insensitive), helix-turn-helix if containing HTH, phage if containing phage (case-insensitive), retroviral integrase if containing rve, and CRISPR-Cas if containing Cas or CRISPR, otherwise domains were assigned to other.

### *V. cholerae* superintegron repeat analysis

To explore the relationship between targets and gene cassettes, anchor-targets were queried in NCBI nucleotide databases, each anchor-target's best E-value identified, hits of the anchor-target having this E-value retained, and the number of such hits to each accession with this E-value computed (Methods). Anchor-targets 1, 4, and 30 had 13, 13, and 10 mappings with 100% query cover and identity each to a single respective assembly, the highest numbers of perfect mappings to single assemblies of any anchor-target having > 0.1% of the total anchor count (Suppl. File 3C).

The 56 targets accounting for >1% of total anchor counts were detected in 223/282 samples (Suppl. File 3B). Of these 56 targets, 27 contain the start codon ATG and 7 contain the stop codon TAG, suggesting that targets, extending beyond the end of the *Vibrio cholerae* repeat, may be transcribed and constitute the 5' or 3' UTRs or regulatory signals for cassette genes.

Anchor-target 1 had 13 perfect mappings to *V. cholerae* chr2 (AP023370.1); 1/13 contained a hypothetical protein in the adjacent inter-repeat region; 12/13 contained no predicted protein annotation (Suppl. File 3B). Further, there was a predicted ORF but no predicted gene annotation in 10/13 perfect mappings (AP023370.1) (Suppl. File 3B). In each of anchor-target 1's 13 perfect mappings, the inter-repeat sequence was extracted from the assembly; we generated a consensus sequence using the 13 extracted inter-repeat sequences (Suppl. File 3D, 3E). The consensus BLASTx-aligned in frame -3 to disulfide interchange protein DsbD (e-values of  5e-63, 5e-51, and 7e-59) and to DUF3709 (e-value of 2e-59). Multiple sequence alignment of these 4 database proteins to the 13 inter-repeat sequences, translated in frame -3, showed high coverage and identity with the database proteins (Supplemental File 3F, Supplemental Figure 6). Anchor-target 4 had 13 100% cover and identity hits to *V. mimicus* chr 1 (CP077425.1) and 10 such hits to *V. cholerae* chr 2 (CP053803.1). Anchor-target 30 had 10 100% cover and identity hits to *V. cholerae* chr 2 (CP077198.1). In each such alignment, the adjacent inter-repeat region contained a predicted DUF645 cassette gene.To explore the high number of DUF645 copies, the superintegron array was approximated as the region between the left- and right-most appearances of the anchor in the two analyzed assemblies to which anchor-targets 4 and 30 had the most perfect mappings. GenBank flat files were extracted from these approximated regions, thus reporting the assemblies' predicted genes in approximate superintegron array; this does not count predicted genes with internal stops or frameshifts (Suppl. File 3G, 3H).

To further investigate the prediction that proteins with DUF645 were enriched in copy number, we analyzed repeat regions of genome assemblies from *V. cholerae* chr 2 (CP077198.1): the most prevalent annotations were of hypothetical proteins (47 copies, respectively), but the second-most prevalent predicted proteins were in the DUF645 family, with 38 copies out of 204 total predicted proteins in the repeat regions (Suppl. File 3G, 3H). Multiple sequence alignment of the DUF645 family cassette genes extracted from GenBank flat files in *V. cholerae* chr 2 (CP077198.1) showed the protein sequences were structurally varied but generally homologous, supporting the hypothesis that a single cassette gene has proliferated in the superintegron array.

**Analysis of a putative CRISPR repeat**

Among the 395 anchors in Table 3 failing BLASTN alignment (Table 3, Suppl. File 2A), multiple sequence alignment of the 7th most highly-entropic BLAST-unaligned anchor's compactors displayed a pattern resembling a CRISPR repeat array: the anchor is contained in a 36-bp element repeating at regular intervals; distinct sequences interspace this repeat (Suppl. File 2C. None of the 23 read-supported compactors aligned using BLASTx, and none of the 10 most supported anchor-targets aligned using BLASTN. Three compactors BLASTN-aligned to MAGs annotated as phage, with e-values of 1e-04 (BK040081.1), 0.016 (OP031041.1), and 0.005 (BK018822.1) (Suppl. File 2D).

FASTQs from PacBio (ERR4025905) and Oxford Nanopore (ERR6713714, ERR6713710, ERR6712627) long-read sequences were found using Pebblescout search of metagenomics databases and searched using SRA BLAST for the repetitive 36mer, which aligned to 9 reads (Suppl. File 2E, 2F).

The first BLASTx hit to a Cas protein was investigated for each read. In 2/9 reads, the best BLASTx hits were not to Cas, but to serine hydroxymethyltransferase (e-value 9e-55, protein accession: MDO5386537.1, read gnl|SRA|ERR6713714.114503.1) and to A/G-specific adenine glycolase (e-value 2e-44, protein accession MBP3515979.1, read gnl|SRA|ERR6712627.32917.1). All reads aligned to one of Cas1, Cas2, or Cas9 with e-values ranging from 2e-125 to 1e-14 (Suppl. File 2G). Despite these e-values, amino-acid identity between the translated nucleotide sequences and the database proteins in the aligned frame was as low as 412/1149 (35.8%) and 244/615 (39.7%), indicating the proteins are diverged (Suppl. Figure 3B, Suppl. File 2G). Only 1,576 amino acids of the 3,762 total covered by alignment had shared identity across the 9 reads, and individual reads had average percent identity of 49.8%.

**Analysis of Anchors in Driving *V. cholerae* Sample Clusters**

In Figure 5E, anchors are classified as ICE-aligning if at least one of anchor-targets 1 and 2 had a BLASTN alignment containing the substring "ICE."

In the anchor having the most post-2018 counts (1076.0, AAATGATGGTCTGAAACAAACGGTAGT), anchor-targets 1 and 2 both megaBLAST-aligned to *V. cholerae* SXT element genomic sequence (KT151663.1). They align to a hypothetical protein (ALP45200.1) which, aligned with BLASTp, has 100% identity and cover to WYL-domain containing protein in *Gammaproteobacteria* (WP_000180271.1) and 100% cover and 94% identity to WYL-domain containing protein in *V. cholerae* (EGQ9107866.1). The two single-nucleotide variations between anchor-targets 1 and 2 would constitute codon changes: GTG to GGG and CCA to ACA (Val to Gly, Pro to Thr).

In another anchor among top 5 post-2018 counts (1065.0, TTTTAACACCTTAGGTATTAATATACC), anchor-targets 1 and 2 both megaBLAST-aligned to *V. cholerae* SXT element genomic sequence (KT141663.1). Anchor-target 1 mapped with 100% cover and identity. These anchor-targets, distinguished by 1 SNV, did not appear in a predicted protein region, but were between predicted genes annotated as ynv (ALP45175.1) and traA (conjugative transfer protein, ALP45176.1).

The anchor having the highest pre-2019 counts was CTAAGCTTCTTCAGCCAACCGTTCTGC, which also megaBLAST-aligned to *V. cholerae* SXT element genomic sequence (KT141663.1). Only anchor-target 1 did so, but it did with 100% query cover and identity, 57 basepairs upstream of the 2 previously discussed sequences. This suggests anchor-target 2 should have been aligned here, as it differs from anchor-target 1 by only 1 base pair. This alignment is within the predicted protein ynv (ALP45175.1). BLASTp of the protein sequence shows it has 274/277 identical amino acids and 100% cover to putative MosA antitoxin component, partial [*Enterovibrio nigricans*] (CCD10491.1). 51 / 100 BLASTp results, all with e-value 0, were to type IV toxin-antitoxin system AbiEi family antitoxin.
ACG -> GCG Thr -> Ala. Both anchor-targets 1 and 2 aligned to *Proteus mirabilis* strain Ire01 integrative and conjugative element SXT/R391 genomic sequence (MN520463.1), and appeared within a predicted protein transcriptional regulator AbiEi 3 antitoxin Type IV TA system (QKR72262.1).

**Filters Used to Select Anchors in Table 1**

| Dataset | Effect Size Threshold | Entropy Threshold | # Anchors Processed |
|---|---|---|---|
| Group A *Streptococcus* (Kachroo et al. 2019) | 0.9 | 0.9 | 8043 |
| Capsule Intestinal Profiling (Shalon et al. 2023) | 0.99 | 0.99 | 8252 |
| *E. coli* (GenomeTrakr, PRJNA230969) | 0.95 | 0.95 | 7560 |
| *Streptococcus pneumoniae* (Chaguze et al. 2020) | 0.9 | 0.9 | 7198 |
| IBDMDB | 0.99 | 0.99 | 8117 |
| Group B *Streptococcus* | 0.99 | 0.99 | 7840 |

| | | | |
|---|---|---|---|
| (Chaguza et al. 2022) | | | |
| Group A *Streptococcus* (Southon et al. 2020) | 0.9 | 0.9 | 7930 |
| *V. cholerae* (LeGault et al. 2021) | 0.9 | 0.9 | 3429 |
| *L. monocytogenes* (Maury et al. 2016) | 0.9 | 0.9 | 7200 |
| *M. tb* lineage 2 (CRypTIC) | 0.9 | 0.9 | 1516 |
| *M. tb* lineage 3 (CRyPTIC) | 0.9 | 0.9 | 1210 |

**Anchor Preprocessing in Figure 2B**

Anchors are reported if failing Bowtie2 alignment to databases representing *C. idella*: GCF_019924925.1 (grass_carp), *A. aegypti*: GCF_002204515.2_AeagL5.0 (aedes), and *Z. marina*: GCA_001185155.1_Zosma_marina.v.2.1(zostera_marina).

6. Krueger, F. *TrimGalore: A wrapper around Cutadapt and FastQC to consistently apply adapter and quality trimming to FastQ files, with extra functionality for RRBS data*. (Github).
7. Leplae, R., Hebrant, A., Wodak, S. J. & Toussaint, A. ACLAME: a CLAssification of Mobile genetic Elements. *Nucleic Acids Res.* **32**, D45–9 (2004).
8. Albertin, C. B. *et al.* The octopus genome and the evolution of cephalopod neural and morphological novelties. *Nature* **524**, 220–224 (2015).
9. Kalvari, I. *et al.* Non-Coding RNA Analysis Using the Rfam Database. *Curr. Protoc. Bioinformatics* **62**, e51 (2018).
10. Berardini, T. Z. *et al.* The Arabidopsis information resource: Making and mining the 'gold standard' annotated reference plant genome. *Genesis* **53**, 474–485 (2015).
11. Ross, K. *et al.* TnCentral: a Prokaryotic Transposable Element Database and Web Portal for Transposon Analysis. *MBio* **12**, e0206021 (2021).
12. The UniVec database. http://www.ncbi.nlm.nih.gov/tools/vecscreen/univec/.
